# Differences between suspended and sinking particles regulate carbon flux in the upper mesopelagic during a *Phaeocystis* Bloom

**DOI:** 10.64898/2026.06.09.731151

**Authors:** Carolina Cisternas-Novoa, Elisa Romanelli, Uta Passow

## Abstract

Despite decades of research, the factors determining the sinking velocity of marine biogenic particles remain poorly constrained, and growing evidence suggests that particle composition and morphology are as important as size in determining particle fate. We compared characteristics of suspended and sinking particles at three depths below the mixed layer and within the layer of maximal flux attenuation during the decline of a *Phaeocystis pouchetii* bloom in the Labrador Sea using marine snow catchers.

Biochemical and morphological characteristics of suspended and sinking particles always differed, with differences depending primarily on bloom stage, and depth accounting for comparatively less variation. Exopolymer particles played a key role, with the relative concentrations of transparent exopolymer particles consistently higher in the suspended than in the sinking particle fraction. In contrast, the partitioning of coomassie-stainable particles changed with the bloom stage, as a function of the *Phaeocystis* life cycle. Ballast minerals played a negligible role during the late-bloom and bloom-decline stages, and their relative importance increased during the non-bloom stage. The C:N ratio was lower in suspended than sinking particles, with differences in morphological measures depending on bloom stage.

Our findings emphasize that export potential is driven not only by particle size, but also by bloom stage, which is closely linked to plankton community composition and plays a key role in the timing and magnitude of carbon flux in the upper mesopelagic. Further, this work highlights the important and diverse roles of exopolymers in regulating carbon flux.

## Introduction

The Biological Carbon Pump transfers organic carbon from the surface to the deep ocean, where it can be stored for decades to centuries (Volk and Hoffert 1985; Ducklow et al. 2001). The efficiency of the gravitational settling pathway, the primary vehicle of carbon transport of the pump (Boyd et al. 2019; Le Moigne 2019; Nowicki et al. 2022; Siegel et al. 2023), depends on the balance between the sinking velocity and the loss rate of sinking particles. Measuring particle sinking velocity in the ocean is not simple, and we rely on estimates of this parameter. Despite the widespread use of size-based parameterizations in biogeochemical models, accumulating evidence shows that internal particle characteristics, such as porosity and composition, frequently reflect sinking potential more accurately than size (Iversen and Lampitt 2020; Omand et al. 2020; Cael et al. 2021; Williams and Giering 2022). However, comprehensive characteristics-based analyses of natural marine particles, especially those comparing suspended and sinking particles across ecological gradients, remain scarce.

Sediment traps and *in situ* pumps have provided insights into the composition of sinking and suspended particles, respectively (Wakeham and Lee 1993; Abramson et al. 2010; Rontani et al. 2011); however, differences in temporal and spatial scales across such methods prevent direct comparisons. The marine snow catcher (MSC) addresses this gap by separating sinking from suspended particles within the same water mass, enabling the simultaneous sampling of both particle types (Riley et al. 2012; Giering et al. 2016; Romanelli et al. 2024). Thus, MSC data allow investigation of how biogeochemical composition and morphological characteristics shape particle sinking behavior.

While the role of mineral ballast in increasing excess density, sinking velocity, and carbon export has been studied (Armstrong et al. 2002; Klaas and Archer 2002; Fischer and Karakaş 2009; Cram et al. 2018), the impact of exopolymeric particles on sinking velocity is just starting to be investigated (Romanelli et al. 2023; Yamada et al. 2024). Transparent exopolymeric particles (TEP), which are rich in acidic polysaccharides (Alldredge et al. 1993), enhance aggregation (Alldredge et al. 1993; Passow 2002) but are intrinsically buoyant unless ballasted (Alldredge and Crocker 1995; Azetsu-Scott and Passow 2004; Mari et al. 2017). The sinking velocities of marine snow depend, among other things, on the relative concentrations of TEP (Engel and Schartau 1999; Azetsu-Scott and Passow 2004; Chajwa et al. 2024). Coomassie stainable particles (CSP) that contain protein (Long and Azam 1996) are also ubiquitous and have been detected in the deep ocean (Cisternas-Novoa et al. 2015; Nagata et al. 2021), but their role in aggregation and export remains poorly characterized.

The Labrador Sea is a hotspot for carbon export, accounting for approximately 15% of the global export (Sanders et al. 2014). Diatoms have commonly dominated the spring phytoplankton bloom in the Labrador Sea, but large *Phaeocystis pouchetii* blooms dominated in 2015 and 2022 (Devred et al. 2025). *P. pouchetii*, which is extensively distributed in cold waters of the Northern Hemisphere, may exist as single-celled, motile flagellates or form large mucous-rich colonies with hundreds of cells each (Verity et al. 1988; Schoemann et al. 2005; Smith and Trimborn 2024). During large blooms, the colonial stage is ubiquitous. As *Phaeocystis* blooms senesce, colonies lose structural integrity and rupture (van Boekel et al. 1992). The fate of the carbon produced during these large *Phaeocystis* blooms varies; although substantial sedimentation can occur, most of the exported material is retained in the upper mesopelagic rather than reaching deeper waters (Reigstad and Wassmann 2007; Smith et al. 2017). Consequently, the contribution of *Phaeocystis*-derived carbon to long-term carbon sequestration is considered relatively limited (Reigstad and Wassmann 2007; Wolf et al. 2016; Roca-Marti et al. 2025; Laget et al. 2025).

In May 2022, during the Biological Carbon Export in the Labrador Sea (BELAS-1) program, an anomalously large, spatially heterogeneous *Phaeocystis pouchetii* bloom was observed in the Labrador Sea (Devred et al. 2025). Primary production during this and a similarly unusual *P. pouchetii* bloom in 2015 was exceptionally high compared to other years (Devred et al. 2025). However, the fate of the carbon generated during these abnormal blooms is unclear (Devred et al. 2025). During the waning of the 2022 *P. pouchetii* bloom, we investigated how biogeochemical and morphological differences between suspended and sinking particles shape carbon export, with a special focus on the role of exopolymeric particles. We hypothesized (1) that inking particles consistently differ from suspended ones, and (2) that the characteristics of both particle types vary with the bloom stage and depth.

## Methods

### Study Site and Sample Collection

During the BELAS-1 expedition on board the R/V Celtic Explorer, we collected suspended and sinking particles in the Labrador Sea (57-59.5° N/ 49-50° W) from May 19^th^ to June 2^nd^, 2022, using MSCs (Fig. 1a). MSCs were deployed at eight stations, at three depths each. As water for all analysis could not be sampled from every MSC, samples for exopolymers or biominerals were collected only in half of the deployments. Sampling stations were categorized into one of three *bloom stages*, allowing us to combine results from different stations.

**Figure 1:**
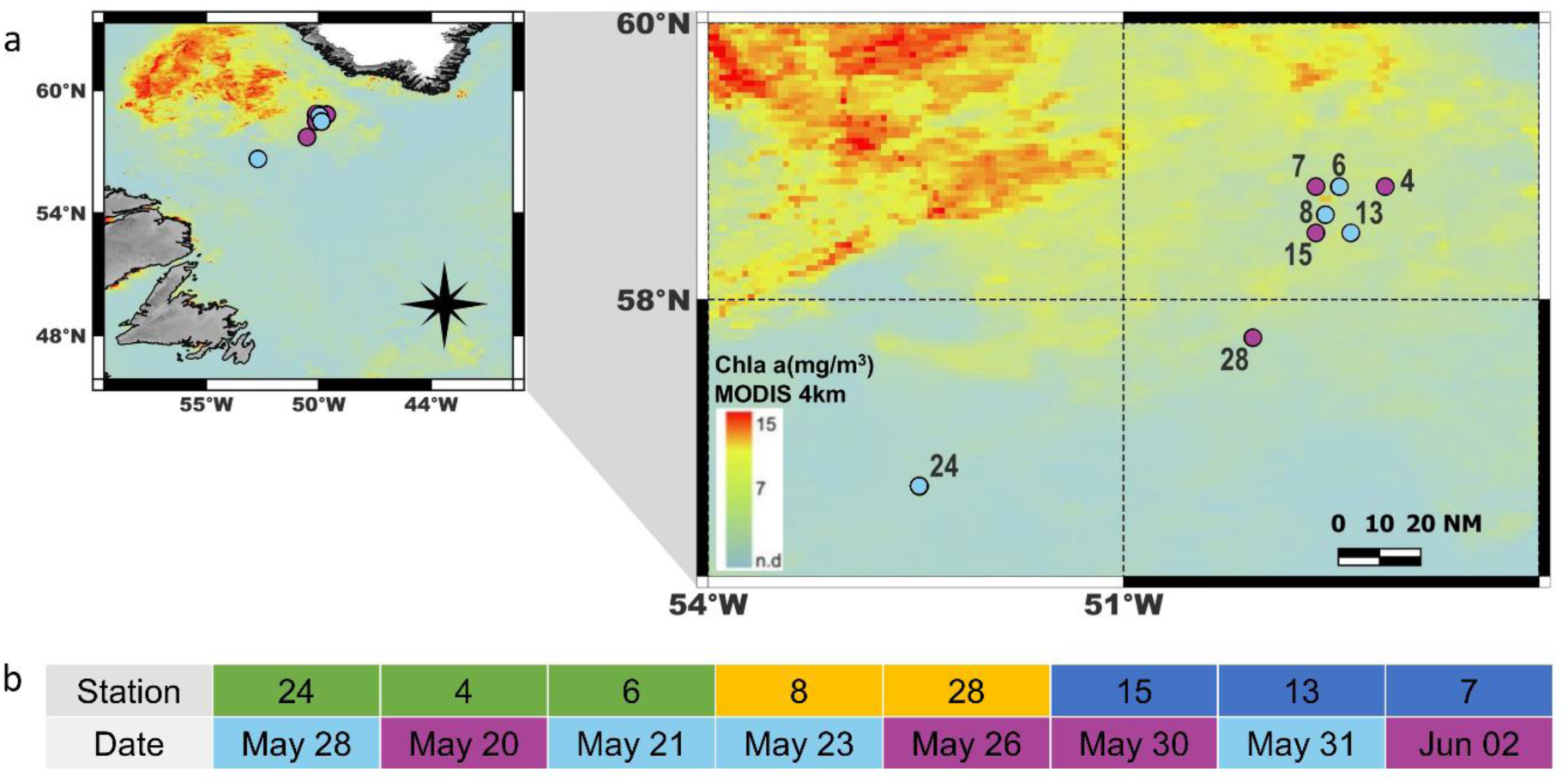
Study area, showing the eight stations sampled with MSCs. (a) Cyan colored markers indicate stations where samples for exopolymers were collected, while magenta marks stations where biomineral samples were taken. The background depicts a MODIS Chl.-a map, reflecting a four-kilometer monthly mean during the sampling period from May to June 2022. (b) Station numbers are shown, colored according to their bloom stage designation (green = late-bloom; yellow = bloom-decline; blue = non-bloom), and their sampling dates are colored based on whether samples were collected for exopolymers or biominerals.

Before each MSC deployment, vertical water column profiles were obtained using the SBE 911 CTD unit with an SBE 32 carousel. CTD profiles were used to identify the mixed-layer depth (between 15 and 77 m; Table 1) as the depth where potential density differed by 0.03 kg m⁻³ from the surface (Brainerd and Gregg 1995). The calibrated fluorescence sensor was used to estimate chlorophyll a (Chl. -a) concentrations. The mixed layer depth was always shallower than the primary productive depth (Table 1), at which the fluorescence reached 10% of its maximum value (Owens et al. 2015).

**Table 1.** Marine Snow Catcher (MSC) deployment depths, D1 to D3, primary production zone (PPZ), and mixed-layer depth (MLD) at each station.

| Stage | Station | MSC deployment depth (m) |  |  | PPZ (m) | MLD (m) |
| --- | --- | --- | --- | --- | --- | --- |
|  |  | D1 | D2 | D3 |  |  |
| Late-bloom | St. 24 | 60 | 160 | 300 | 118 | 53 |
|  | St. 4 | 80 | 125 | 300 | 138 | 15 |
|  | St. 6 | 80 | 125 | 300 | 151 | 20 |
| Bloom-decline | St. 8 | 150 | 250 | 500 | 182 | 39 |
|  | St. 28 | 120 | 220 | 500 | 159 | 65 |
| Non-bloom | St. 15 | 90 | 190 | 300 | 110 | 41 |
|  | St. 13 | 120 | 220 | 500 | 192 | 77 |
|  | St. 7 | 70 | 170 | 500 | 65 | 29 |

### Marine Snow Catchers Sampling

The 150 cm-tall MSC consists of a 100-liter water sampler with a removable base section (7.6 L), which facilitates separation of suspended and sinking particles based on differences in sinking velocity (Riley et al. 2012; Giering et al. 2016). Particles were harvested in three operationally-defined fractions following a 2-hour settling period (Romanelli et al. 2023, 2024). Suspended particles were collected from the central tap located ∼79 cm from the top of the MSC (top fraction), corresponding to particles with sinking velocity considered negligible in terms of carbon export and sequestration (≤ ∼9 m d⁻¹). Slow-sinking particles, which had maximal sinking velocities under 18 m d⁻¹, were siphoned from the upper part of the base section. Fast-sinking particles, defined by velocities exceeding 18 m d⁻¹, were collected in a tray at the bottom of the base section. Before subsampling the tray, photographs of its contents were taken using a 48 MP digital camera with optical image stabilization to quantify later sinking marine snow (> 0.1mm) concentrations and sizes.

We collected samples at three depths in the zone of maximal particulate organic carbon (POC) flux attenuation (twilight zone), where remineralization is most intense (e.g., Buesseler et al. 2007; Boyd and Trull 2007): The shallowest MSC, located below the MLD (60 m to 150 m) and within the primary productive zone will be called D1. D2 is defined as the transition depth between 125 m and 260 m, generally below the productive zone, and D3 was located in the upper mesopelagic between 300 m and 500 m (Table 1). For most MSC deployments, pressure sensors (RBR Duet 3) were installed to verify deployment depths.

We collected samples to analyze particulate organic carbon (POC) and particulate nitrogen (PN) at all stations. 50 mL subsamples, fixed with formalin (1%-2% final concentration), were collected for FlowCam analysis at all stations except stations 24 and 13. Samples for TEP and CSP analysis were collected at stations 24, 6 (late-bloom), 8 (bloom-decline), and 13 (non-bloom), whereas samples for biogenic silica (BSi) and particulate inorganic carbon (PIC) were taken at stations 4 (late-bloom), 28 (bloom-decline), 15, and 7 (non-bloom; Fig. 1b). Thus, each analysis was conducted at least once per depth per bloom stage.

Concentrations of particles from the fast-sinking pool were determined by subtracting the base section from the tray, while the concentrations of slow-sinking particles were obtained by subtracting the top section from the base section and correcting for volume (Riley et al. 2012; Giering et al. 2016) (Table S1). For data analysis, slow-sinking and fast-sinking particles were merged and collectively referred to as the “sinking fraction”, which was compared to suspended particles.

### CTD Sampling for Chlorophyll a and Phaeopigments

Vertical profiles of Chl.-a were derived from CTD-mounted fluorometer measurements (SBE 911 CTD) and corrected using high-pressure liquid chromatography (HPLC) derived Chl.-a concentrations. HPLC sampling was conducted at 10 depths, from the surface to 200 m, with sample volumes from 0.5 to 2.2 L. The samples were filtered through 25mm Whatman GF/F filters and stored at −80°C until HPLC analysis for pigments (Claustre et al. 2004). Chl.-a and phaeopigments (Pheo) at depth D1 will be used to define the bloom stages.

### Chemical Analysis

#### Total Particulate Carbon, Particulate Organic Carbon, and Particulate Nitrogen

We determined the concentrations of total particulate carbon, POC, and particulate nitrogen (PN) in the top, base, and tray of the MSC by filtering, in duplicate, 1 L, 0.75 L, and 0.2 L, respectively, through pre-combusted (450°C, 6 h) glass fiber filters (25 mm diameter GF/F, Whatman). Filters were dried (40°C, 12h) onboard and stored in a desiccator until analysis. After acid fuming (10% HCl) of POC/PN filters, all filters were analyzed using a PerkinElmer 2400 CHN Analyzer with acetanilide standards. The accepted deviation of elemental analysis results is 0.4%. The blank values were C = 28 ± 1 mg and N = 0.49 ± 0.2 mg. We calculated PIC by subtracting POC from the total particulate carbon measured without prior acidification. C:N ratios are molar.

#### Transparent exopolymer particles and coomassie stainable particles

TEP and CSP were measured spectrophotometrically following Passow and Alldredge (1995) and Cisternas-Novoa et al. (2014), respectively, with TEP calibration based on Bittar et al. (2018). On board, we filtered two replicates of each MSC fraction (0.1–1 L) onto 0.4 μm polycarbonate filters (25 mm, Nuclepore, Whatman). Filters were stained immediately with 0.02% Alcian Blue (pH 2.5, Sigma Aldrich) or 0.04% Coomassie Brilliant Blue G-250 (pH 7.4, SERVA electrophoresis), for TEP and CSP, respectively, and then stored frozen (–20 °C) until analysis. TEP concentrations are expressed as micrograms of xanthan gum equivalents per liter (μg XG eq. L⁻¹), and CSP concentrations are expressed as micrograms of bovine serum albumin equivalents per liter (μg BSA eq. L⁻¹).

#### Biogenic Silica

We determined biogenic silica (BSi) concentrations by filtering 0.1 to 1 L of the sample through polycarbonate membrane filters with a 0.6-µm pore size and a 47 mm diameter (Isopore, Millipore). The filters were dried at 40°C for 12 hours and stored in a desiccator until analysis. For digestion, we placed the filters in Teflon tubes, added 4 mL of 0.2 N NaOH, heated at 95°C for 40 minutes, and then cooled. To measure BSi, 4 mL of the solution was diluted with 6 mL of Milli-Q water and analyzed using the molybdosilicic acid spectrophotometric method (Strickland and Parsons 1968).

### Particle Imaging

#### Imaging of Particles < 1mm

Using a FlowCam 8000 Series (Fluid Imaging, Inc.; Sieracki et al. 1998; Camoying and Yñiguez 2016), we assessed the morphological characteristics and size distribution of small particles (diameter < 1mm) at each station from formalin-fixed samples that were first filtered through a 1 mm mesh. We conducted all measurements in automatic imaging mode at 2× magnification, using flow cells with a depth of 1000 µm, which enabled us to measure particles ranging from 6 µm to 1000 µm in diameter. We analyzed sample volumes ranging from 0.5 mL to 2 mL at a flow rate of 3–4 mL min⁻¹, with a camera rate of 7.0 frames s⁻¹. Between samples, we flushed the system with Milli-Q water for 5–10 minutes at a flow rate of 0.5 mL min⁻¹ and visually inspected the flow cell to ensure cleanliness. During each run, we gently mixed samples with a small stirring bar to prevent particle sedimentation and pumped them through the flow cell with a high-precision syringe pump.

We used FlowCam Visual Spreadsheet© software version 5.8.12 (Fluid Imaging, Inc.) to analyze particle size, shape, and other morphological and optical properties. Particle size was quantified as area-based diameter, calculated from a circle with an area equal to the area of the particle’s edge trace. We binned particles into octave-scale size classes (65–130 µm, 130–260 µm, 260–520 µm, 520–1040 µm, and >1040 µm) starting at 65 µm, where a linear trend emerged to estimate the differential number size distributions N(D) (# particles mL⁻¹ mm⁻¹) of the suspended and sinking fractions. We calculated the slope by fitting a one-degree polynomial to the log10-transformed particle size distribution and log10-transformed arithmetic mean of each size bin (μm). In addition to size, we analyzed irregularity (FlowCam’s “compactness”), particle transparency, and a proxy for particle porosity: P*_int_*. Irregularity is calculated based on the particle’s perimeter and filled area; values > 1 indicate more convoluted outlines, while a perfect circle has a value of 1. We use the term irregularity rather than “compactness” to avoid confusion with its standard meaning: the quality of being closely packed. Particle transparency quantifies shape solidity, with values near 1 for filled, circular shapes and lower values for elongated, irregular, or porous forms. P_int_ (intensity normalized by size) is also a proxy for particle porosity (Bach et al. 2019). It was calculated from intensity, which refers to the mean grayscale value per pixel or brightness; higher values indicate greater light transmission, which is typical in porous aggregates with low solid content (Yokogawa Fluid Imaging Technologies, Inc. 2023). Irregularity and porosity allow, for example, the differentiation between aggregates and solid primary particles like cells.

#### Marine Snow Imaging

All photographs of marine snow collected in the MSC trays reflect sinking marine snow > 0.1mm without artifacts from handling, staining, or fixation, which may destroy colony integrity (Schoemann et al. 2005; Wassmann et al. 2005; Smith and Trimborn 2024). These photographs of sinking marine snow were visually inspected to classify concentrations as high, moderate, or low and size the marine snow as large, medium, or small. At D1, photographs were additionally analyzed in ImageJ to enumerate and size sinking marine snow and estimate total volume (in ppm) using the equivalent spherical diameter. To characterize the particle size distribution of sinking marine snow, the equivalent spherical diameters were binned into log₁₀-spaced classes (0.1–20 mm), and bins with fewer than two particles were excluded. A linear regression was fitted to the log–log relationship between equivalent spherical diameter and abundance. The slope of this fit, along with its standard error and 95% confidence interval, describes the particle-size distribution of sinking marine snow, providing insights into the relative contribution of large vs. small particles within the analyzed size range.

#### Imaging of TEP and CSP

For select samples, TEP and CSP were analyzed microscopically using the methods by Alldredge et al. (1993) and Long and Azam (1998). On board, 10 to 100 ml of the sample was filtered through 0.4 µm polycarbonate filters and stained with Alcian Blue (TEP) or Coomassie Brilliant Blue (CSP). Filters were mounted on the white side of semi-transparent glass slides (CytoClear, Poretics Corp., Livermore, US; Logan et al., 1994) and stored frozen at –20 °C until analysis. Images for qualitative assessment were captured with a light microscope (Olympus CX41RF) equipped with a camera (Lumenera 2-1R 1.4 Megapixel CCD Series).

#### Calculation of Instantaneous POC Flux Attenuation Coefficient (*b*)

To assess an instantaneous POC flux attenuation, we calculated the attenuation coefficient (*b*) from MSC profiles at the three depths, using the power-law model (Martin et al. 1987):

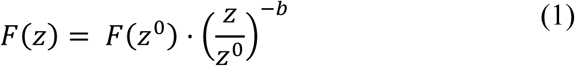

where F(z) is the flux at depth z, and z₀ is a fixed reference depth (80 m). The model was linearized by log-transforming both flux and normalized depth, and *b* was calculated as the negative slope of the linear regression of log-transformed POC flux against log(z/z₀). The coefficient of determination (R²) was used to assess the goodness of fit. Note that this *b* value of instantaneous flux attenuation is not directly comparable with *b* values based on annual fluxes as measured, for example, by moored traps.

POC fluxes of slow-sinking (F_slow_) and fast-sinking (F_fast_) particles at each depth were calculated to provide the basis for *b* estimates:

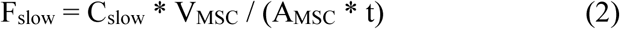

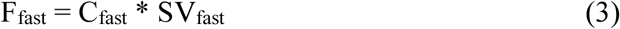

where C_slow_ and C_fast_ are the POC concentrations of slow and fast sinking particles, respectively, V_MSC_ is the volume of the MSC (89.9 L including base), A_MSC_ is the area of the MSC base (0.06 m^2^), t is the settling time (2 h), and SV_fast_ denotes the mean sinking velocity of the fast-sinking fraction. We estimated a conservative lower bound of flux (i.e., F_slow_ + F_fast_), assuming that the concentration gradient observed in the MSC formed within 2 hours. This likely underestimates the true flux (Giering et al., 2016).

## Statistical Analysis

Generally, the quantitative data are presented as the mean (average) ± standard deviation, first averaging over replicates and then averaging between stations within each bloom stage (POC, TEP, CSP, marine snow abundance), with error propagation. Morphological particle characteristics are given as means with the 95% confidence intervals. Standard errors reflect the uncertainty of the PSD slopes. To compare the biochemical composition of suspended and sinking particles, exopolymer and biomineral concentrations were normalized to POC (Table S2).

We tested several variables across the three bloom stages using a one-way ANOVA, followed by Tukey’s post hoc test to identify significant pairwise differences (p < 0.05). Tested variables included marine snow concentration, slope of the marine snow particle size distribution, POC, and C:N ratio in the sinking fraction, and Chl.-a concentration, the pheopigment-to-Chl.-a ratio (Pheo/Chl.-a) in the water column, all at D1, as well as the instantaneous POC flux attenuation.

The biochemical composition of suspended and sinking particles was compared using a Wilcoxon rank-sum test to assess differences between both particle types when data were pooled across all bloom stages and depths. We selected the non-parametric Wilcoxon test because some variables were not normally distributed, and sample sizes were relatively small. We considered the results statistically significant at p < 0.01.

Differences in particle size and morphology (Irregularity, Transparency, and P_int_) between bloom stages were tested using the Kruskal–Wallis test followed by Dunn’s post-hoc tests with Benjamini-Hochberg correction. Significance was accepted at p < 0.001.

A Factor Analysis of Mixed Data (FAMD), which included 10 quantitative and qualitative variables (Table S3), explored relationships among biogeochemical, morphological, and ecological variables across depths and bloom stages. This method integrates qualitative variables, such as bloom stage, depth, and particle type, as well as, the quantitative variables representing biochemical composition and morphological characteristics. We conducted the analysis using the package FactoMineR in R (Lê et al. 2008) and retained five dimensions based on eigenvalue thresholds and the proportion of variance explained. Despite the modest sample size, the FAMD effectively integrated mixed data types and revealed coherent multivariate structure across bloom stages and particle types.

## Results

To investigate how sinking particles differ from suspended ones and to determine whether most of the *P. pouchetii* carbon sank in the upper mesopelagic, samples collected using the marine snow catchers were analyzed in relation to bloom stage and depth. Classification of stations into different bloom stages was necessary because of the spatially patchy progression of the bloom (Devred et al. 2025).

### Characterization of the Bloom Stages

Each station sampled with the MSC was associated with one of three bloom stages based on statistically significant differences in characteristics of sinking particles below the mixed layer at D1 (Table S4). Specifically, bloom stages were designated based on (i) the concentrations of sinking marine snow, (ii) the slope of the associated marine snow size distributions, (iii) the POC concentrations in the sinking fraction, (iv) the water column Chl.-a concentration, and (v) the associated Pheo/Chl.-*a* ratio, all at D1, as well as on (vi) the instantaneous POC flux attenuation coefficient (*b*). Supporting evidence, albeit statistically not significant (Table S4), was provided by the C:N ratio of sinking particles. The three identified bloom stages are hereafter referred to as late-bloom, bloom-decline, and non-bloom.

The late-bloom stage was characterized by high concentrations of sinking POC and marine snow, and high Chl.-a in the water column (Fig. 2), and the least steep slope of the marine snow size distribution (−1.63 ± 0.16), indicating a high proportion of large, sinking marine snow at D1. The instantaneous POC flux attenuation coefficient was highest at the late bloom stage (*b* = 1.52 ± 0.29; Table S4), revealing that POC flux declined strongly with depth. The molar C:N ratio of sinking particles at D1 was relatively low (7.3 ± 1.4) compared with the following bloom stages. The Pheo/Chl.-a ratio was also low, indicating limited Chl.-a degradation and thus the presence of fresher material in the water column (Table S4).

**Figure 2.**
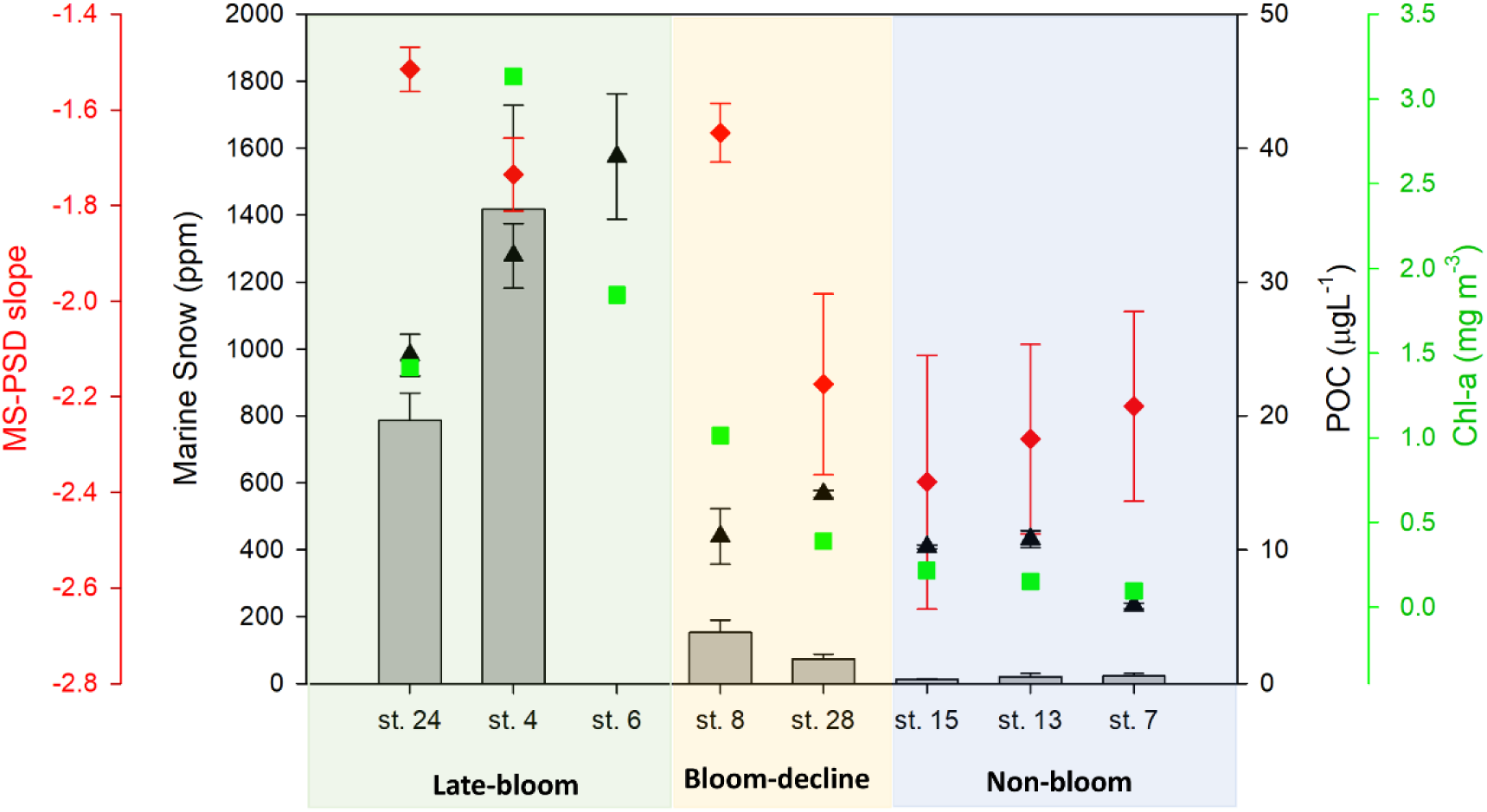
Definition of bloom stages: Marine snow concentrations (grey bars), slopes of marine snow size distributions (MS-PSD slope, red diamonds), POC concentration of the sinking fraction (black triangles), and total Chl.-a concentration in the water column (green squares) at D1, reveal significant differences between the three bloom stages. Error bars represent standard deviations (marine snow, POC) or standard errors (MS-PSD slope).

At bloom-decline, concentrations of sinking POC, Chl.-a, and marine snow at D1 were significantly lower compared to the late-bloom stage (Fig. 2), and the slope of the particle size distribution of sinking marine snow was steeper (−1.91 ± 0.37), indicating a shift toward smaller marine snow. The lower *b*-value (0.67 ± 0.22) indicates reduced attenuation of POC flux at this time. The C:N ratio (9.1 ± 2.8) was elevated, indicating less fresh material than during late-bloom. The high Pheo/Chl.-a ratio supports increased degradation as the bloom transitions into senescence.

During the non-bloom stage, sinking POC, Chl.-a, and marine snow were at their lowest concentrations compared to the other two stages (Fig. 2). The slope of the size distribution of sinking marine snow was steepest at this stage (−2.29 ± 0.08), indicating a dominance of small marine snow. The *b*-value of 0.53 ± 0.02 (Table S4) suggests that although biomass was low, instantaneous flux attenuation was comparably low. The elevated C:N ratio (9.8 ± 4.3) relative to the other bloom stages, together with the high Pheo/Chl.-a ratios (0.76 ± 0.40), confirms a greater degradation state of organic matter compared to late-bloom (Table S4).

Several other features differed across the bloom stages, including the morphological characteristics of particles ranging from 6 to 1000 µm and the microbial community composition. When averaged over depth, suspended and sinking particles showed significant differences in particle size (ABD), porosity (P_int_), irregularity, and transparency between bloom stages, except for the transparency of the suspended particles, which did not differ significantly between late-bloom and non-bloom (Table S5). The microbial community in the water column as queried via 16S rRNA gene metabarcoding also differed significantly between the bloom stages: While *P. pouchetii* dominated the photosynthetic community (> 95% of chloroplast 16S rRNA) during all three stages, the relative abundance of the photosynthetic community compared to the non-photosynthetic bacterial ASV dropped from > 50% to 7-23% to < 5%, respectively, from the late-bloom, to the bloom-decline to the non-bloom stage. *Thalassiosira spp.* (diatom) was the second most prevalent photosynthetic genus and occurred at a higher relative abundance (up to 4%) at the non-bloom stage compared to the late-bloom and bloom-decline stages (both < 1%) (Romanelli et al. 2026). Furthermore, underwater vision profiler (UVP) results confirm the presence of intact, large *P. pouchetii* colonies (4-6 L^-1^) at 40-80 m during late-bloom and bloom-decline, but negligible concentrations of such colonies at the non-bloom stage (Laget et al. 2025). Our classification of stations into bloom stages provides an environmental context for assessing differences in particle composition and morphology as a function of ecosystem state and is not meant to represent a chronological continuum of a single phytoplankton bloom.

### Suspended and sinking particles

Consistent with previous MSC studies (Riley et al., 2012; Giering et al., 2016; Baker et al., 2017; Romanelli et al., 2023, 2024) most particles belonged to the suspended rather than the sinking particle fraction, with the suspended fractions of POC (>78%), PN (>60%), TEP (>85%), CSP (>66%), and BSi (>59%) consistently exceeding the sinking fractions across all bloom stages and depths (Fig. S1). The only exception was PIC during the non-bloom stage at D3, where sinking particles dominated PIC concentrations (66%). The large fraction of suspended particles, even at mesopelagic depth (D3), emphasizes that fragmentation of sinking marine snow must play a central role at depth (Briggs et al. 2020; Siegel et al. 2025).

Independent of bloom stage, suspended particles always differed significantly from sinking ones in their C:N and TEP/POC ratios (Table S6). Mean C:N ratios of suspended particles were lower than those of sinking particles (5.2 ± 0.4 vs. 8.8 ± 2.2), whereas mean relative TEP concentrations were higher in suspended than in sinking particles (2.3 ± 2.0 vs. 0.1 ± 0.1 XG eq. (µg-C)⁻¹; Table S2). For all other parameters, suspended–sinking differences varied with bloom stage and are examined separately for each stage.

### Partitioning of exopolymer particles between suspended and sinking fractions

TEP/POC ratios were consistently higher in suspended than in sinking particles, with one exception. In contrast, the partitioning of CSP between suspended and sinking particles varied with bloom stage.

During the late-bloom TEP/POC in suspended particles was higher than in sinking particles, except at D3 when relative TEP concentrations were high in both fractions. In contrast, CSP/POC ratios were lower in suspended than in sinking particles at all depths (Fig. 3a, b).

**Figure 3.**
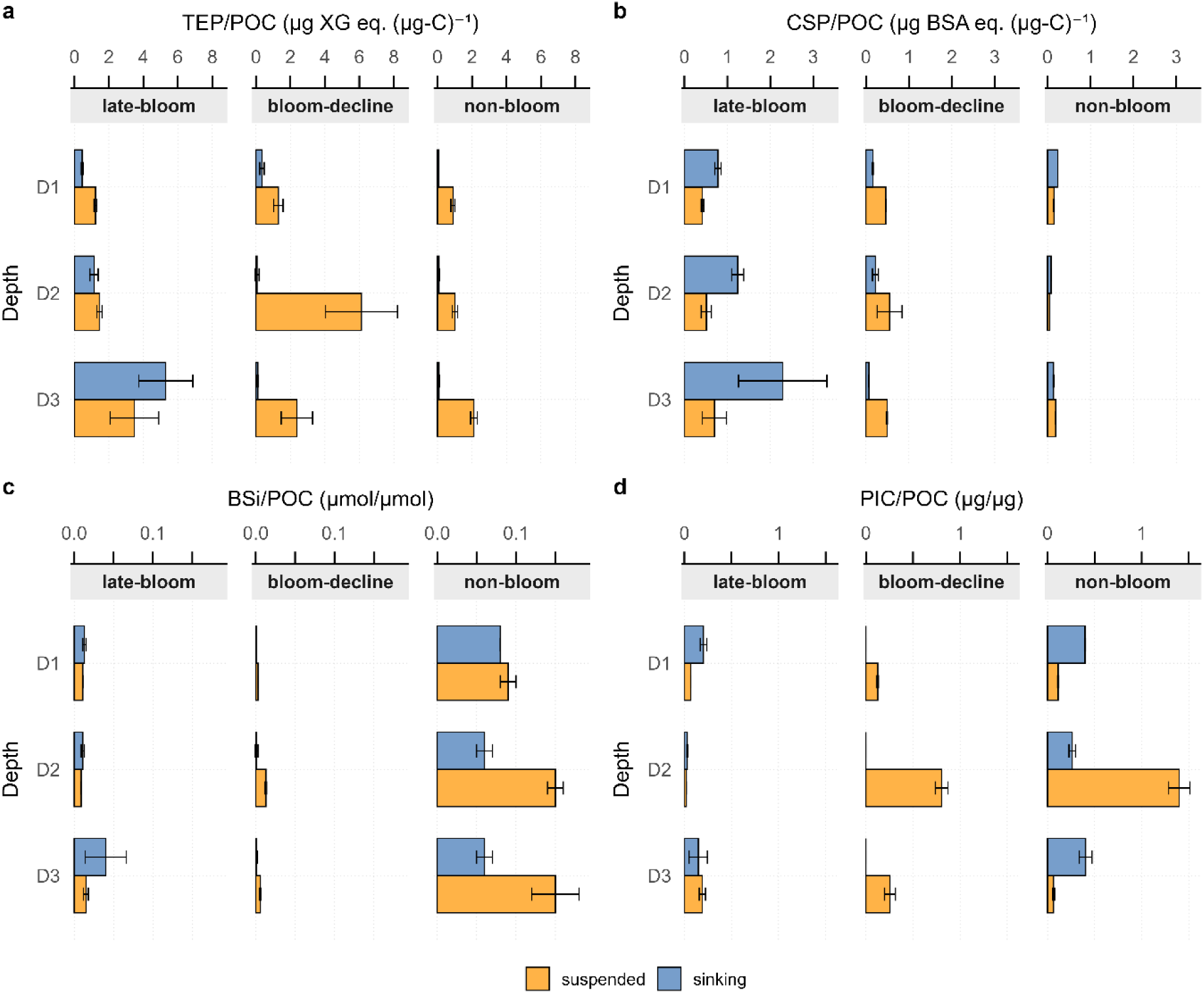
Composition of suspended (yellow) and sinking (blue) particles at each bloom stage and depth: (a) POC normalized transparent exopolymeric particles (TEP/POC in µg XG eq.(µg C)^-1^); (b) POC normalized Coomassie stainable particles (CSP/POC in µg BSA eq.(µg C)^-1^); (c) POC normalized biogenic silica (BSi/POC in µmol Si/µmol C); (d) POC normalized inorganic carbon (PIC/POC µg/µg).

During the bloom-decline stage, the TEP/POC ratios of suspended particles were elevated relative to those in the other bloom stages (Fig. 3a), consistent with expectations of increased colony disruption and TEP release during colony senescence (Verity et al. 1988). The CSP/POC ratios were similar to the TEP/POC ratios and appreciably higher in suspended than in sinking particles at all depths, in contrast to the late-bloom observations (Fig. 3b).

During non-bloom, the relative contribution of TEP was still noteworthy, although lower than during late bloom and bloom-decline, whereas the relative contribution of CSP was very small, and no clear difference in the CSP/POC ratio between suspended and sinking particles was discernible (Fig. 3a, b). High concentrations of TEP are known to persist after *P. pouchetii* blooms, and large amounts of foam may accumulate at the sea surface after such blooms (Lancelot et al. 1987; Smith and Trimborn 2024).

### Partitioning of biominerals between suspended and sinking fractions

During the late-bloom, both the BSi/POC and the PIC/POC ratios were relatively low, as expected during a bloom dominated by *Phaeocystis*. The BSi/POC ratios of suspended and sinking particles were similar in D1 and D2 and higher in sinking particles at D3 (Fig. 3c). No differences existed between particle types in the PIC/POC ratios (Fig. 3d).

During the bloom-decline, the BSi/POC ratios were low and without a notable trend. The PIC/POC ratio of suspended particles was higher than during the late-bloom stage and appreciably higher than that of sinking particles (0.1 to 0.8 μg/ μg; Fig. 3c, d). PIC/POC values > 0.6 are common during blooms of *Emiliania huxleyi* (Findlay et al. 2011), a coccolithophorid known to co-occur with other phytoplankton (Hopkins et al. 2015). However, coccolithophorids were not detected in the 16S rRNA dataset (Romanelli et al. 2026).

During the non-bloom stage, the BSi/POC ratios of both suspended and sinking particles were high compared to the other bloom stages (Fig. 3c), reflecting the greater contribution of diatoms to both suspended and sinking particles (Romanelli et al. 2026). The PIC/POC ratios, especially in sinking particles, were also higher than in the other two stages, suggesting a more diverse eukaryotic microbial community during the non-bloom stage than during earlier bloom stages (Romanelli et al. 2026).

Concentrations of lithogenic particles, which can act as ballast, were not determined in this study. However, aerosol deposition, the primary source of lithogenic material in the central Labrador Sea, far from river or glacial sources, is expected to be low (Menzel Barraqueta et al. 2018). During a follow-up expedition, two years later (2024), low LSi/POC ratios were observed for suspended particles and for sinking particles (0.005 ± 0.005 µmol Si (µmol C)^-1^ and 0.007 ± 0.005 µmol Si (µmol C)^-1^, respectively), suggesting that lithogenic silica likely played a minor role in particle export during the study period.

### Size and morphology of suspended and sinking particles < 1 mm

In late-bloom, at D1, suspended particles were significantly smaller, as well as less porous, and less irregular than sinking ones (Fig. 4; ABD: 16.4 µm vs. 24.9 µm; *P_int_*: 5.2 vs. 8.4; irregularity: 1.02 vs. 1.28). The steeper PSD slope of suspended vs. sinking particles (Table S7; – 2.90 vs. –1.96) confirmed a high proportion of relatively large sinking particles at D1. Large, porous, and irregular particles in the sinking fraction suggest the presence of small sinking marine snow < 1 mm at D1. No differences in size, porosity, or irregularity were detected between suspended and sinking particles at D2 or D3.

**Figure 4.**
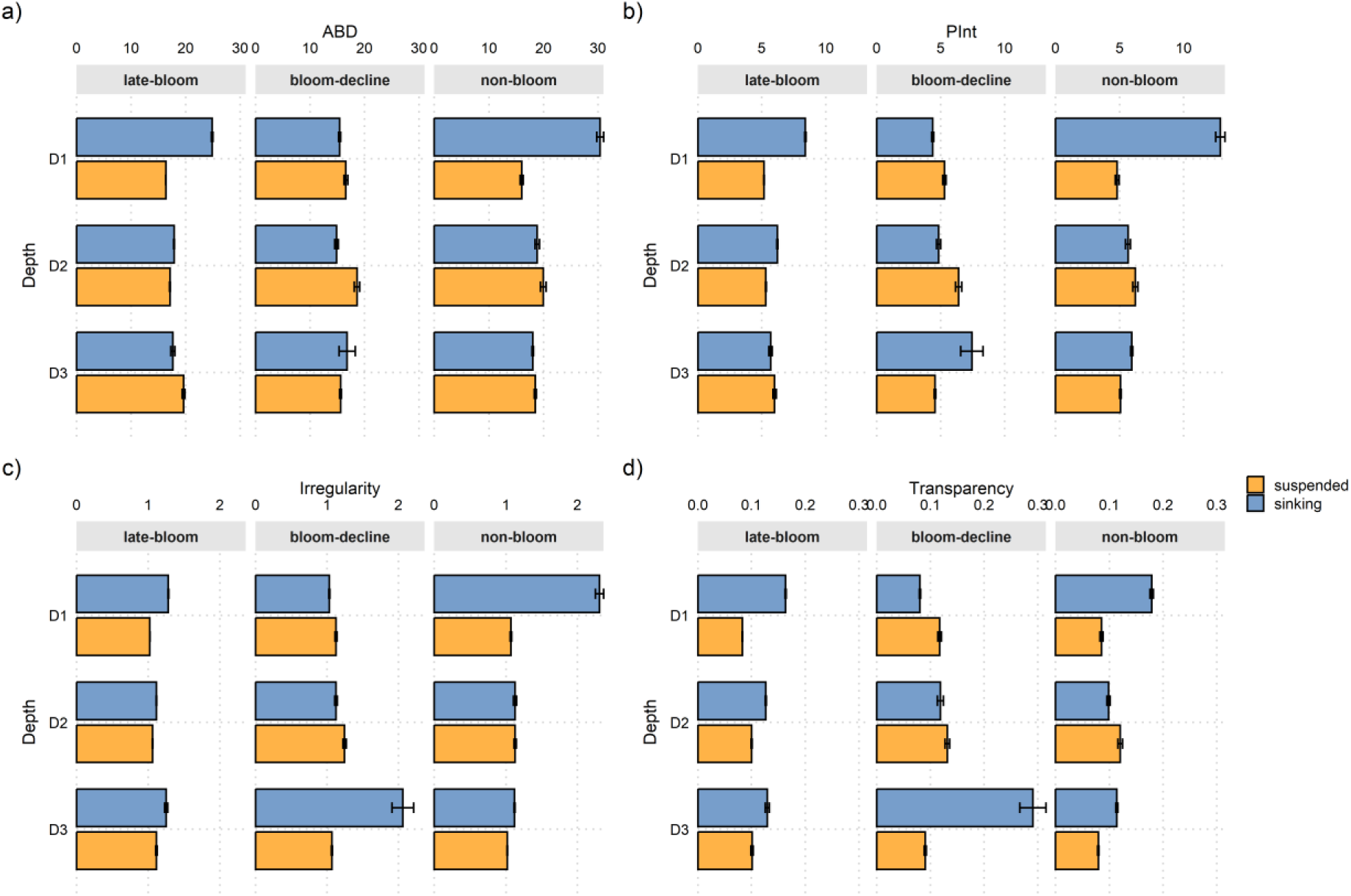
Morphological and optical characteristics of suspended and sinking particles < 1mm: (a) Area-based diameter (ABD, µm), (b) porosity, P_int_, (c) irregularity, and (d) transparency across bloom-stages and depth, given as mean with error bars representing the 95% confidence intervals. The lack of visible error bars means they are too small to see.

During bloom-decline, mean particle sizes of sinking particles were smaller compared to those observed during late-bloom (Fig. 4a). Suspended particles were of similar or larger size than sinking ones, implying low concentrations of small sinking marine snow < 1mm. The slightly higher porosity (Fig. 4b and d) and irregularity (Fig. 4c) of suspended particles compared to sinking ones at D1 and D2, possibly reflects the relatively high fraction of exopolymers in the suspended fraction, especially at D2 (Fig. 3). At D3, in contrast, sinking particles were more porous (*P_int_* 7.44 vs. 6.40) and irregular (2.06 vs 1.25) compared to suspended ones, suggesting that remnants of small, sinking *Phaeocystis* snow dominated sinking particles < 1 mm at this depth.

During non-bloom, suspended particles at D1 were approximately half the size of sinking particles (ABD: 16 µm vs. 30.4 µm). Sinking particles at D1were also larger than those observed during late-bloom and bloom-decline stages (24.9 µm and 15.5 µm, respectively). Furthermore, sinking particles at D1 exhibited more than two-fold greater porosity and irregularity than suspended particles (e.g., *P_in_*_t_: 12.9 vs. 4.8; irregularity: 2.31 vs. 1.07). The high porosity and irregularity, combined with the relatively low exopolymer concentrations and the increased contributions of ballasting biominerals (Fig. 3), suggest the presence of small miscellaneous aggregates < 1mm in size.

### Characteristics of sinking marine snow > 0.1 mm

During the late-bloom stage, large sinking marine snow up to 9 mm in size, which consisted mostly of *Phaeocystis* snow, was present at high concentrations (6416 ± 2181 L^-1^ or 763 ±155 ppm) at D1 (Fig. 2, Table S8). This large sinking *Phaeocystis* snow, which appeared greenish to brownish (Table S9), had lost its intact membrane and most of the colony matrix, but was rich in *Phaeocystis* cells. Samples stained for TEP and CSP revealed that these individual *Phaeocystis* cells were coated with a layer of CSP (Fig. 5), explaining the elevated CSP contribution in sinking particles. Microscopic observations revealed that at D1 and D2, TEP were mostly unattached, which explains their TEP partitioning into the suspended particle fraction. Visual inspection of marine snow at D2 and D3 during the late-bloom showed decreasing concentrations of large sinking marine snow with depth (Table S9, Fig. S2), consistent with the patterns observed for smaller marine snow discussed above.

**Figure 5.**
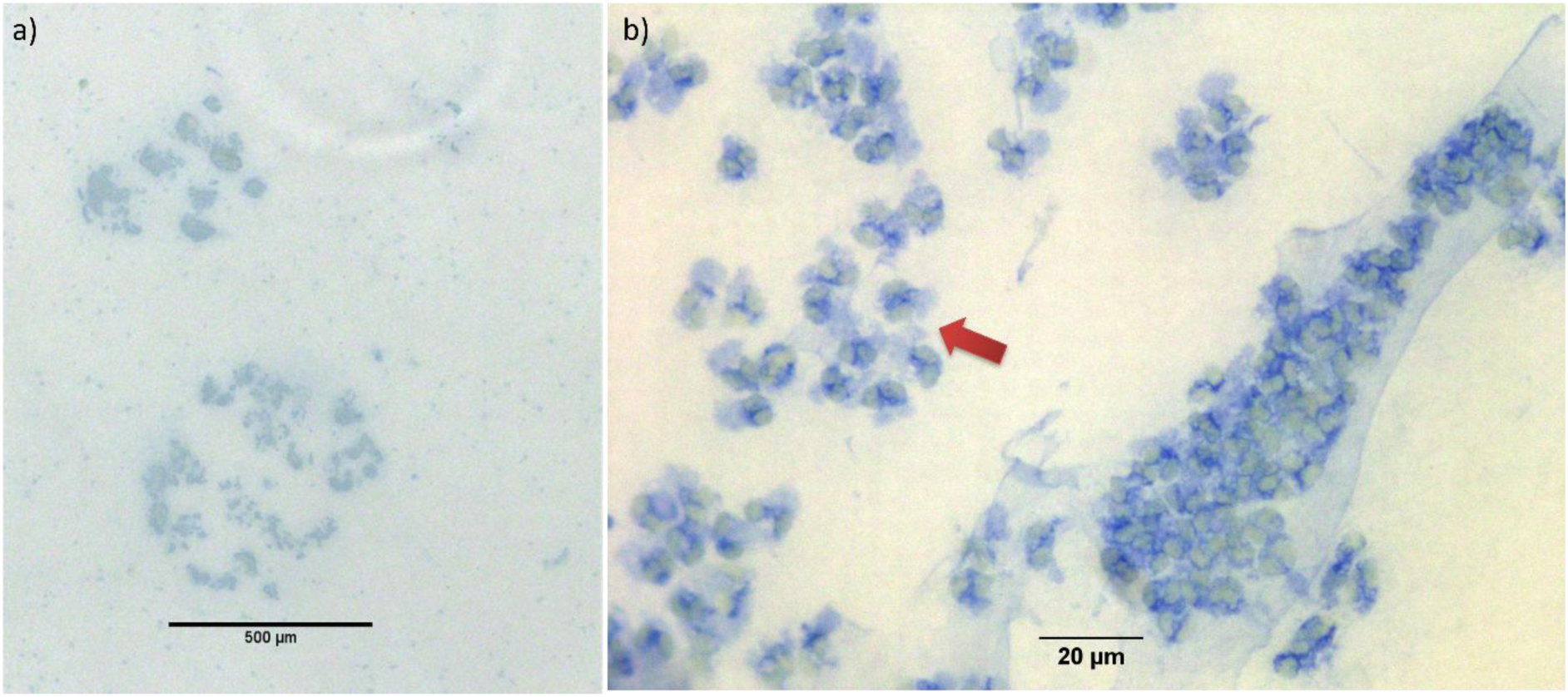
*Phaeocystis pouchetii* snow formed from disintegrating colonies. Photos from the sinking fraction during the late-bloom, stained with Coomassie Brilliant Blue, highlighting the enrichment in CSP. (a) scale bar: 500 µm, (b) arrow indicates CSP acting as a glue, keeping the cells together after colony disruption. Scale bar: 20 µm. Note: While this image was taken after filtering and staining the sample to make CSP visible, the abundance and size of *P. pouchetii* snow are based on photos taken directly in the MSC tray.

During bloom-decline, the concentrations of large, sinking *Phaeocystis* snow at D1 were lower (4320 ± 4965 L^-1^), and its maximum size was smaller, up to 4 mm, compared to the late-bloom stage (Table S8, Fig. S3). As during late-bloom, the abundance and size of large, sinking marine snow decreased with depth (Table S9, Fig. S2). Although macroscopically similar, microscopy revealed that the sinking *Phaeocystis* snow collected at bloom-decline primarily consisted of detrital material with only a few embedded *Phaeocystis* cells, distinguishing it from the *Phaeocystis* snow observed during the late-bloom. The increased contribution of CSP in suspended particles (Fig. 3b) during bloom-decline is consistent with the low abundance of Phaeocystis cells in the sinking fraction. The low concentrations of large sinking *Phaeocystis* snow > 0.1 mm at D3 (Fig. S2) combined with the high porosity and irregularity of small sinking marine snow < 1 mm at that depth (Fig. 4), suggest that *Phaeocystis* snow had continued to disintegrate, forming particles < 0.1 mm, but retaining its high porosity and irregularity.

At the non-bloom stations, at D1, mean concentrations (1397 ± 168 L^-1^; Table S8) and maximum sizes of large, sinking marine snow > 0.1 mm were smaller compared to the other bloom stages (Fig. S2, Fig. S3, Table S8). This contrasts with the findings of small, sinking aggregates < 0.1 mm (Fig. 4), which were relatively large, porous, and irregular at D1, possibly reflecting the presence of small miscellaneous aggregates. Concentrations of such sinking aggregates > 0.1 mm remained relatively constant with depth (Table S9).

### Factor analysis of mixed data (FAMD) of particle characteristics and bloom stages

Results from the Factor Analysis of Mixed Data (FAMD, Chavent et al., 2022) depict distinct differences between suspended and sinking particles and between bloom stages (Fig. 6). While the first five dimensions accounted for 80.3% of the total variance, Dimensions 1 and 2 explained 28% and 19%, respectively, and Dimensions 3, 4, and 5 contributed 14%, 11%, and 9% (Table S10). Dimension 1 primarily captured differences in particle morphology, with the highest contributions from P_int_ (0.80), irregularity (0.78), area-based diameter (0.61), and the slope of the particle size distribution (0.60), followed by a moderate contribution from transparency (0.49). Dimension 2 was driven by CSP/POC (0.71), the C:N ratio (0.47), and the categorical variable bloom stage (0.47), with TEP/POC contributing moderately (0.30).

**Figure 6.**
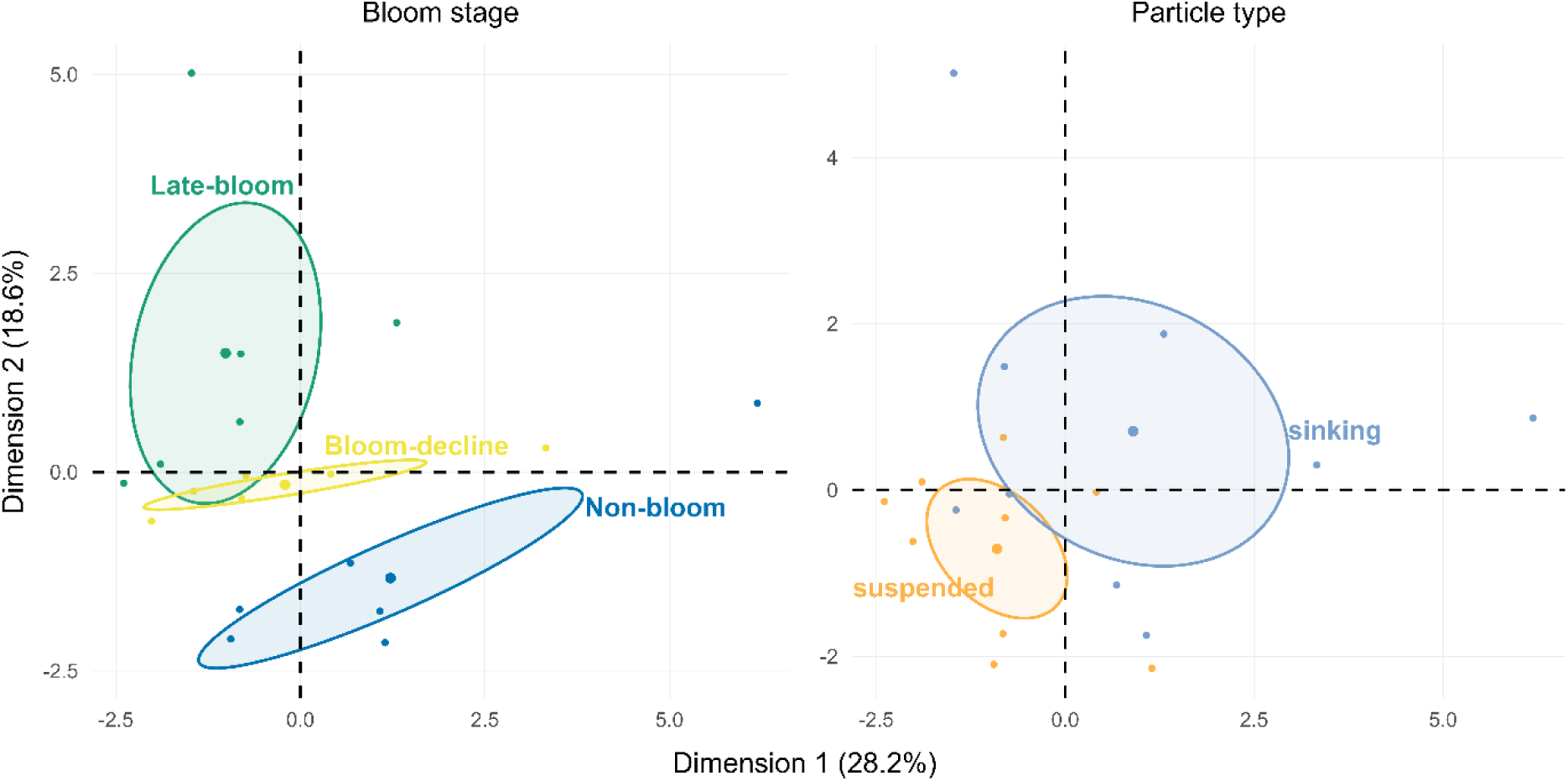
Factor Analysis of Mixed Data (FAMD) plots illustrating particle characteristics across bloom stage (left) and particle types (right). Ellipses represent within-group variance and are included for visual reference only; they do not imply statistical significance or discrete clustering.

The FAMD results (Fig. 6) support our hypotheses that sinking particles differ consistently from suspended particles and that the characteristics of both particle types vary with bloom stage. Particle composition and characteristics clustered by bloom stage, validating our bloom-stage classification based on D1 parameters and indicating that these patterns extend into the upper mesopelagic. In contrast, depth contributed mainly to dimensions 4 and 5, suggesting a weaker influence on particle properties.

## Discussion

### *P. pouchetii* bloom dynamics drive differences between suspended and sinking particles in the upper mesopelagic

During the BELAS-1 expedition in spring 2022, we encountered a large, spatially heterogeneous *P. pouchetii* bloom in its declining phase (Devred et al. 2025). All MSC stations were classified into three specific bloom stages based on particle characteristics below the mixed layer depth, with the classification further supported by plankton community data above the mixed layer. The FAMD, which integrated biochemical and morphological characteristics of suspended and sinking particles, corroborated our hypothesis that particle characteristics below the mixed layer, within the zone of high flux attenuation, differed consistently between suspended and co-located sinking particles as a function of the bloom stage. Detailed analysis revealed that the life cycle of *P. pouchetii,* the dominant primary producer, played a key role in driving these differences between particle types.

The linkages between bloom dynamics and particle properties are schematically illustrated in Fig. 7. During the late-bloom stage, when intact *P. pouchetii* colonies were still abundant above the mixed layer (Laget et al. 2025), *Phaeocystis* snow appeared below the mixed layer (this study). As the protective membrane surrounding *P. pouchetii* colonies (Hamm et al. 1999) became porous and disintegrated, the colony matrix was released into the water as TEP, as has also been observed previously (Lancelot et al. 1987; Passow et al. 1994; Rousseau et al. 2007; Verity et al. 2007; Smith et al. 2017). The buoyancy of TEP (Azetsu-Scott and Passow 2004; Mari et al. 2005, 2017) ensures that the majority of TEP is partitioned into the suspended particle fraction. This sequence of events explains the observed TEP distribution at the different bloom stages (Fig. 7). We discovered that CSP dynamics differed from those of TEP. The fresh sinking *Phaeocystis* snow was rich in CSP-coated cells, which seemed to stick together (Fig. 5). Thus, while TEP was mainly observed in the suspended fraction, CSP was primarily associated with sinking particles during the late-bloom stage (Fig 7a). Instantaneous flux attenuation was high, indicating that the large size of this fresh *Phaeocystis* snow did not translate into high sinking velocities. A relatively high degradation rate of suspended total organic carbon likely contributed further to the high instantaneous flux attenuation during the late-bloom stage (Romanelli et al. 2026).

**Figure 7.**
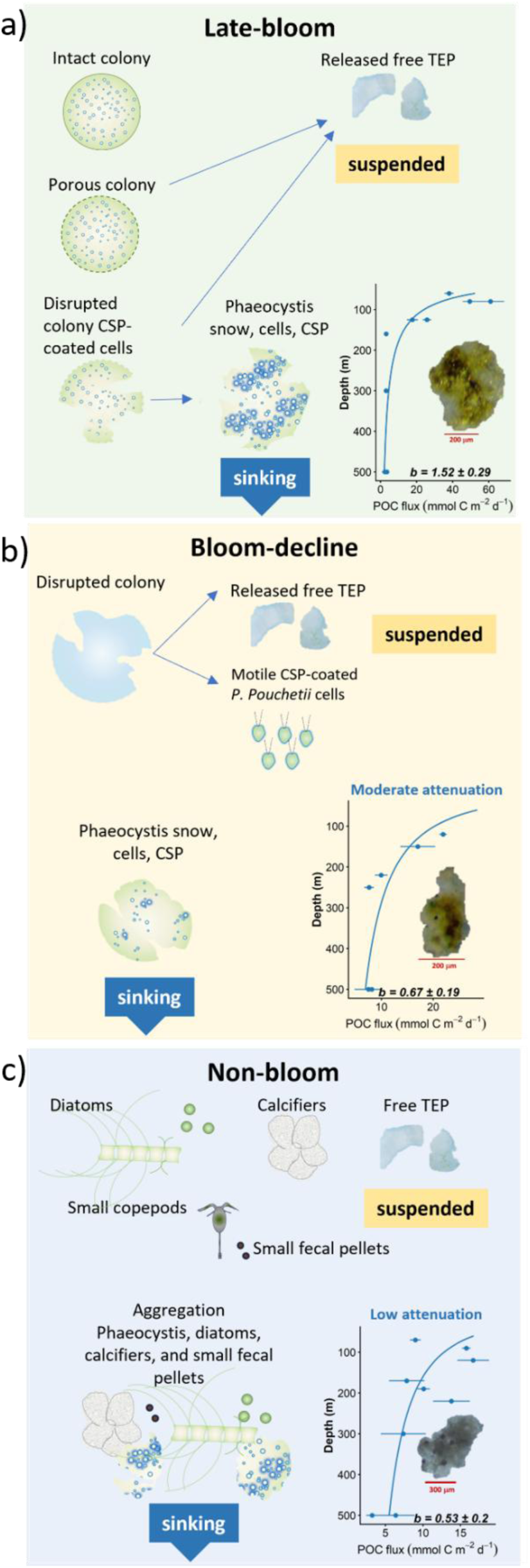
Conceptual model of particle characteristics and sinking behavior during the three stages of a declining *Phaeocystis pouchetii* bloom. The inset in the bottom-right corner shows instantaneous POC flux profiles with power-law regression curves, indicating the estimated *b*-values (± standard error). The reference depth is z_0_ = 80 m. Each curve is accompanied by a representative aggregate image.

During bloom-decline, both CSP and TEP were most abundant in the suspended particle fraction. This may reflect the lifecycle of *Phaeocystis*, where the complete disruption of colonies during bloom collapse allows non-motile, CSP-coated *Phaeocystis* cells to develop flagella, become motile, and emigrate from the colonies (Verity et al. 1988), joining the suspended particle pool (Fig. 7b). Sinking particles were comparatively small, dense, and contained few exopolymer particles compared to late-bloom conditions. These smaller, more uniform particle sizes of sinking particles likely reflect the continued disintegration and degradation of *Phaeocystis* snow. The organic carbon degradation rates of sinking particles were of the same order of magnitude as those of the late-bloom stage (Romanelli et al. 2026). The lower *b-*value combined with similar degradation rates implies that during bloom-decline, mean sinking velocities of the smaller and less porous sinking snow were faster compared to the sinking snow of the late-bloom.

During the non-bloom stage (Fig 7c), concentrations of primary producers were very low; *Phaeocystis* occurred only in their single-cell stage (no colonies), and diatoms such as *Thalassiosira* spp. (Romanelli et al. 2026), as well as calcifiers and small fecal pellets (Laget, pers. comm.) were present. Sinking particles were porous and irregularly shaped, suggesting aggregates formed by the coagulation of various types of particles. These small sinking aggregates contained only small amounts of exopolymeric particles and a relatively high amount of ballast material, typically associated with increased sinking potential (Armstrong et al. 2002; Klaas and Archer 2002; Passow 2004). These particles likely promoted the formation of miscellaneous aggregates through coagulation rather than colony disintegration. These miscellaneous aggregates sank effectively, albeit at low concentrations, due to the overall low biomass. As a result, flux attenuation was low and transfer efficiency relatively high compared to the other bloom stages.

High flux attenuation during senescence of a *Phaeocystis* bloom is consistent with observations that sedimentation of carbon to depths>100 m is generally low during *P. pouchetii* bloom collapse in the North-West Atlantic (reviewed by Reigstad & Wassmann, 2007) and for *Phaeocystis antarctica* in the Southern Ocean (Wolf et al. 2016). The large amounts of TEP released by aging colonies (Passow and Wassmann 1994; Passow 2000) may, however, lead to a post-bloom flux event of carbon produced by *Phaeocystis*: A sedimentation event of amorphous detrital matter and the simultaneous disappearance of TEP from surface waters was linked to a preceding *P. pouchetii* bloom (Riebesell et al. 1995). A bloom of *Phaeocystis antartica* has been reported to have rapidly exported carbon to great depth (up to 500m) in the Ross Sea (DiTullio et al. 2000). The formation of miscellaneous aggregates during the non-bloom stage may represent the beginning phase of such a post-bloom sedimentation event, which could have led to significant *Phaeocystis* carbon export if ballasting particles (diatom frustules, coccolithophorids, dust) became available.

### Ecological controls on particle flux through the upper mesopelagic

This work emphasizes the importance and complexity of exopolymeric particles and their roles in the marine carbon cycle. Whereas TEP are known to be buoyant and remain suspended in the surface waters or accumulate at the sea surface (Wurl and Holmes 2008; Jennings et al. 2017), as well as being essential for aggregate formation and carbon flux by providing the “glue” within marine snow (Alldredge et al. 1993), the role of CSP, which clearly underlie a different dynamic, is much less well understood. Moreover, the roles of these exopolymers have not been systematically considered with respect to the flux of marine snow and marine particles through the mesopelagic.

Our findings also reinforce the idea that export potential is influenced not simply by particle size (Iversen and Lampitt 2020; Omand et al. 2020; Cael et al. 2021; Williams and Giering 2022) but by the biochemical and structural characteristics of particles, which depend on phytoplankton community succession and, hence, bloom dynamics. Here, we show that the life cycle of the dominant phytoplankton determines the characteristics of suspended and sinking particles, thereby influencing flux through the upper mesopelagic zone. During the *P. pouchetii* bloom, exopolymers and their distribution were key in determining flux in the upper mesopelagic. Most bloom-forming diatoms coagulate at bloom termination to form rapidly sinking aggregates that effectively move primary produced carbon to depth (Smetacek 1985). However, only a small fraction of the TEP generated during a diatom bloom is transported within these diatom aggregates that are ballasted by the silica frustules (Passow et al. 2001). Bloom-forming coccolithophorids such as *Emiliania huxleyi* can contribute to the formation of fast-sinking fecal pellets by acting as dense ballast when ingested by zooplankton. (Wal et al. 1995) or aggregate formation, with the latter more likely when bloom demise is due to viral infection (Riebesell et al. 2017). *E. huxleyi* also forms TEP (Harlay et al. 2009), but overall, very little is known about the underlying sedimentation mechanisms (Balch 2018). The role of sinking is also not well understood for *Trichodesmium* (Berman-Frank et al. 2007; Bonnet et al. 2023; Fourquez et al. 2025), a filamentous diazotrophic cyanobacteria that forms large blooms in the tropical and subtropical oceans (Fourquez et al. 2025). *Trichodesmium* spp. generates large amounts of TEP under stress, and upon programmed cell death, the loss of gas vesicles can lead to the sedimentation of a large fraction of the population (Berman-Frank et al. 2007). Transport of the primary produced carbon to 1000 m depth may, however, be less efficient compared to non-filamentous diazotrophs (Bonnet et al. 2023). These examples emphasize that shifts in dominant phytoplankton composition determine particle characteristics with far-reaching consequences for carbon export in the upper mesopelagic, underscoring the importance of incorporating ecological factors into carbon export concepts and models.

## Supporting information

CisternasNovoa et al_supplemental material

## Acknowledgements

This work was supported by the Northwest Atlantic Biological Carbon Pump (NWA-BCP) project, part of the Ocean Frontier Institute, funded by Canada First Research Excellence Fund awards. UP was supported by the Canada Research Chair Program, and UP and ER were supported by NASA Grant 80NSSC17K0692. We sincerely thank the crew and scientific team aboard the RV Celtic Explorer during the BELAS-1 expedition, as well as the entire NWA-BCP team. We also thank Eduard Leymarie and the laboratory of Hervé Claustre for HPLC analyses of pigments and phaeopigments. We have no conflict of interest.

## Author Contributions

CCN contributed to the conceptualization, data acquisition, data curation, formal analysis, investigation, visualization, and writing of the original draft. ER contributed to data acquisition, the investigation, formal analysis, and the revision of the manuscript. UP contributed to the conceptualization, funding acquisition, project administration, supervision, validation, review, and editing of the manuscript. All authors approved the final version of the manuscript.

## Data Availability Statement

The data used in this study are available through the Canadian Integrated Ocean Observing System (CIOOS) Atlantic Data Catalog (https://catalogue.cioosatlantic.ca/dataset/ca-cioos_a3cf18b4-5bae-4ce4-b274-6498db8c8ebc). Metadata and access instructions are provided in the repository.

## References

Abramson, L., C. Lee, Z. Liu, S. G. Wakeham, and J. Szlosek. 2010. Exchange between suspended and sinking particles in the northwest Mediterranean as inferred from the organic composition of in situ pump and sediment trap samples. Limnology and Oceanography 55: 725–739. doi:10.4319/lo.2010.55.2.0725

Alldredge, A. L., and K. M. Crocker. 1995. Why do sinking mucilage aggregates accumulate in the water column? Science of The Total Environment 165: 15–22. doi:10.1016/0048-9697(95)04539-D

Alldredge, A. L., U. Passow, and B. E. Logan. 1993. The abundance and significance of a class of large, transparent organic particles in the ocean. Deep Sea Research Part I: Oceanographic Research Papers 40: 1131–1140. doi:10.1016/0967-0637(93)90129-Q

Armstrong, R., C. Lee, J. Hedges, S. Honjo, and S. Wakeham. 2002. A new, mechanistic model for organic carbon fluxes in the ocean based on the quantitative association of POC with ballast minerals. Deep Sea Research Part II: Topical Studies in Oceanography 49: 219–236. doi:10.1016/S0967-0645(01)00101-1

Azetsu-Scott, K., and U. Passow. 2004. Ascending marine particles: Significance of transparent exopolymer particles (TEP) in the upper ocean. Limnology and Oceanography 49: 741–748. doi:10.4319/lo.2004.49.3.0741

Bach, L. T., P. Stange, J. Taucher, E. P. Achterberg, M. Algueró-Muñiz, H. Horn, M. Esposito, and U. Riebesell. 2019. The Influence of Plankton Community Structure on Sinking Velocity and Remineralization Rate of Marine Aggregates. Global Biogeochemical Cycles 33: 971–994. doi:10.1029/2019GB006256

Baker, C. A., S.A. Henson, E. L. Cavan, S.L.C. Giering, A. Yool, M. Gehlen, A. Belcher, J.S. Riley, H. E. K. Smith, and R. Sanders. 2017. Slow-sinking particulate organic carbon in the Atlantic Ocean: Magnitude, flux, and potential controls. Global Biogeochemical Cycles 31: 1051–1065. doi:10.1002/2017GB005638

Balch, W. M. 2018. The Ecology, Biogeochemistry, and Optical Properties of Coccolithophores. Ann Rev Mar Sci 10: 71–98. doi:10.1146/annurev-marine-121916-063319

Berman-Frank, I., G. Rosenberg, O. Levitan, L. Haramaty, and X. Mari. 2007. Coupling between autocatalytic cell death and transparent exopolymeric particle production in the marine cyanobacterium Trichodesmium. Environ Microbiol 9: 1415–1422. doi:10.1111/j.1462-2920.2007.01257.x

Bittar, T. B., U. Passow, L. Hamaraty, K. D. Bidle, and E. L. Harvey. 2018. An updated method for the calibration of transparent exopolymer particle measurements. Limnology and Oceanography: Methods 16: 621–628. doi:10.1002/lom3.10268

van Boekel, W., F. Hansen, R. Riegman, and R. Bak. 1992. Lysis-induced decline of Phaeocystis spring bloom and coupling with the microbial food web. Marine Ecology-progress Series - MAR ECOL-PROGR SER 81: 269–276. doi:10.3354/meps081269

Bonnet, S., M. Benavides, F.A.C. Le Moigne, M. Camps, A. Torremocha, O. Grosso, C, Dimier, D. Spungin, I. Berman-Frank, L. Garczarek, F.M. Cornejo-Castillo. 2023. Diazotrophs are overlooked contributors to carbon and nitrogen export to the deep ocean. The ISME Journal 17: 47–58. doi:10.1038/s41396-022-01319-3

Boyd, P. W., H. Claustre, M. Levy, D. A. Siegel, and T. Weber. 2019. Multi-faceted particle pumps drive carbon sequestration in the ocean. Nature 568: 327–335. doi:10.1038/s41586-019-1098-2

Boyd, P. W., and T. W. Trull. 2007. Understanding the export of biogenic particles in oceanic waters: Is there consensus? Progress in Oceanography 72: 276–312. 10.1016/j.pocean.2006.10.007

Brainerd, K. E., and M. C. Gregg. 1995. Surface mixed and mixing layer depths. Deep Sea Research Part I: Oceanographic Research 42: 1521–1543. doi:10.1016/0967-0637(95)00068-H

Briggs, N., G. Dall’Olmo, and H. Claustre. 2020. Major role of particle fragmentation in regulating biological sequestration of CO2 by the oceans. Science 367: 791–793. doi:10.1126/science.aay1790

Buesseler, K. O., C. H. Lamborg, P. W. Boyd, P. J. Lam, T. W. Trull, R. R. Bidigare, J. K. B. Bishop, K. L. Casciotti, F. Dehairs, M. Elskens, et al. 2007. Revisiting carbon flux through the ocean’s twilight zone. Science 316: 567–570. 10.1126/science.1137950

Cael, B. B., E. L. Cavan, and G. L. Britten. 2021. Reconciling the Size-Dependence of Marine Particle Sinking Speed. Geophysical Research Letters 48: e2020GL091771. doi:10.1029/2020GL091771

Camoying, M. G., and A. T. Yñiguez. 2016. FlowCAM optimization: Attaining good quality images for higher taxonomic classification resolution of natural phytoplankton samples. Limnology and Oceanography: Methods 14: 305–314. doi:10.1002/lom3.10090

Chajwa, R., E. Flaum, K. D. Bidle, B. Van Mooy, and M. Prakash. 2024. Hidden comet tails of marine snow impede ocean-based carbon sequestration. Science 386: eadl5767. doi:10.1126/science.adl5767

Chavent, M., V. Kuentz-Simonet, A. Labenne, and J. Saracco. 2022. Multivariate Analysis of Mixed Data: The R Package PCAmixdata.doi:10.48550/arXiv.1411.4911

Cisternas-Novoa, C., C. Lee, and A. Engel. 2014. A semi-quantitative spectrophotometric, dye-binding assay for determination of Coomassie Blue stainable particles. Limnology and Oceanography: Methods 12: 604–616. doi:10.4319/lom.2014.12.604

Cisternas-Novoa, C., C. Lee, and A. Engel. 2015. Transparent exopolymer particles (TEP) and Coomassie stainable particles (CSP): Differences between their origin and vertical distributions in the ocean. Marine Chemistry 175: 56–71. doi:10.1016/j.marchem.2015.03.009

Claustre, H., S. B. Hooker, L. Van Heukelem, J.-F. Berthon, R. Barlow, J. Ras, H. Sessions, C. Targa, C. S. Thomas, D. van der Linde, and J.-C. Marty. 2004. An intercomparison of HPLC phytoplankton pigment methods using in situ samples: application to remote sensing and database activities. Marine Chemistry 85: 41–61. doi:10.1016/j.marchem.2003.09.002

Cram, J. A., T. Weber, S. W. Leung, A. M. P. McDonnell, J.-H. Liang, and C. Deutsch. 2018. The Role of Particle Size, Ballast, Temperature, and Oxygen in the Sinking Flux to the Deep Sea. Global Biogeochemical Cycles 32: 858–876. doi:10.1029/2017GB005710

Devred, E., S. Clay, M. Ringuette, T. Perry, M. Amirian, A. Irwin, and Z. Finkel. 2025. Net primary production in the Labrador Sea between 2014 and 2022 derived from ocean colour remote sensing based on ecological regimes. Remote Sensing of Environment 323: 114713. doi:10.1016/j.rse.2025.114713

DiTullio, G. R., J. M. Grebmeier, K. R. Arrigo, M. P. Lizotte, D. H. Robinson, A. Leventer, J. P. Barry, M. L. Van Woert, and R. B. Dunbar. 2000. Rapid and early export of Phaeocystis antarctica blooms in the Ross Sea, Antarctica. Nature 404: 595–598. doi:10.1038/35007061

Ducklow, H., D. Steinberg, and K. Buesseler. 2001. Upper Ocean Carbon Export and the Biological Pump. oceanog 14: 50–58. doi:10.5670/oceanog.2001.06

Engel, A., and M. Schartau. 1999. Influence of transparent exopolymer particles (TEP) on sinking velocity of Nitzschia closterium aggregates. Mar. Ecol. Prog. Ser. 182: 69–76. doi:10.3354/meps182069

Findlay, H. S., P. Calosi, and K. Crawfurd. 2011. Determinants of the PIC: POC response in the coccolithophore Emiliania huxleyi under future ocean acidification scenarios. Limnology and Oceanography 56: 1168–1178. doi:10.4319/lo.2011.56.3.1168

Fischer, G., and G. Karakaş. 2009. Sinking rates and ballast composition of particles in the Atlantic Ocean: implications for the organic carbon fluxes to the deep ocean. Biogeosciences 6: 85–102. doi:10.5194/bg-6-85-2009

Fourquez, M., F-E. Ababou, M. Camps, F. Van Wembeke, O. Grosso, A. Barani, S. Nurige, L. Guyomarch, F.A.C. Le Moigne, and S. Bonnet. 2025. Aggregation and remineralization of Trichodesmium unveil potential for ocean carbon sequestration. ISME Commun 5: ycaf128. doi:10.1093/ismeco/ycaf128

Giering, S. L. C., R. Sanders, A. P. Martin, C. Lindemann, K. O. Möller, C. J. Daniels, D. J. Mayor, and M. A. St John. 2016. High export via small particles before the onset of the North Atlantic spring bloom. Journal of Geophysical Research: Oceans 121: 6929–6945. doi:10.1002/2016JC012048

Hamm, C. E., D. A. Simson, R. Merkel, and V. Smetacek. 1999. Colonies of Phaeocystis globosa are protected by a thin but tough skin. Marine Ecology Progress Series 187: 101–111.

Harlay, J., C. De Bodt, A. Engel, S. Jasen, Q. d’Hoop, J. Piontek, N. Van Oostende, S. Groom, K. Sabbe, and L. Chou. 2009. Abundance and size distribution of transparent exopolymer particles (TEP) in a coccolithophorid bloom in the northern Bay of Biscay. Deep Sea Research Part I: Oceanographic Research Papers 56: 1251–1265. doi:10.1016/j.dsr.2009.01.014

Hopkins, J., S. A. Henson, S. C. Painter, T. Tyrrell, and A. J. Poulton. 2015. Phenological characteristics of global coccolithophore blooms. Global Biogeochemical Cycles 29: 239–253. doi:10.1002/2014GB004919

Iversen, M. H., and R. S. Lampitt. 2020. Size does not matter after all: No evidence for a size-sinking relationship for marine snow. Progress in Oceanography 189: 102445. doi:10.1016/j.pocean.2020.102445

Jennings, M. K., U. Passow, A. S. Wozniak, and D. A. Hansell. 2017. Distribution of transparent exopolymer particles (TEP) across an organic carbon gradient in the western North Atlantic Ocean. Marine Chemistry 190: 1–12. doi:10.1016/j.marchem.2017.01.002

Klaas, C., and D. E. Archer. 2002. Association of sinking organic matter with various types of mineral ballast in the deep sea: Implications for the rain ratio. Global Biogeochemical Cycles 16. doi:10.1029/2001GB001765

Laget, M., A. Rogge, M. Wietz, L. Musselman, Y-H. Hu, N. McGinty, S. Tuo, M. Holtappels, Z. Finkel, and A. Waite. 2025. First in situ imaging of large colonial Phaeocystis quantifies inefficient carbon export.doi:10.21203/rs.3.rs-7773706/v1

Lancelot, C., G. Billen, A. Sournia, T. Weisse, E. Colijn, M. W. J. Veldhuis, A. Davies, and P. Wassman. 1987. Phaeocystis blooms and nutrient enrichment in the continental coastal zones of the North Sea. Ambio 16: 38–46.

Le Moigne, F. A. C. 2019. Pathways of Organic Carbon Downward Transport by the Oceanic Biological Carbon Pump. Frontiers in Marine Science 6: 634. doi:10.3389/fmars.2019.00634

Lê, S., J. Josse, and F. Husson. 2008. FactoMineR: An R Package for Multivariate Analysis. Journal of Statistical Software 25: 1–18. doi:10.18637/jss.v025.i01

Logan, B. E., H.-P. Grossart, and M. Simon. 1994. Direct observation of phytoplankton, TEP and aggregates on polycarbonate filters using brightfield microscopy. J Plankton Res 16: 1811–1815. doi:10.1093/plankt/16.12.1811

Long, R., and F. Azam. 1996. Abundant protein-containing particles in the sea. Aquat. Microb. Ecol. 10: 213–221. doi:10.3354/ame010213

Mari, X., U. Passow, C. Migon, A. B. Burd, and L. Legendre. 2017. Transparent exopolymer particles: Effects on carbon cycling in the ocean. Progress in Oceanography 151: 13–37. doi:10.1016/j.pocean.2016.11.002

Mari, X., F. Rassoulzadegan, C. P. D. Brussaard, and P. Wassmann. 2005. Dynamics of transparent exopolymeric particles (TEP) production by *Phaeocystis globosa* under N- or P-limitation: a controlling factor of the retention/export balance. Harmful Algae 4: 895–914. doi:10.1016/j.hal.2004.12.014

Martin, J. H., G. A. Knauer, D. M. Karl, and W. W. Broenkow. 1987. VERTEX: carbon cycling in the northeast Pacific. Deep Sea Research Part A. Oceanographic Research Papers 34: 267–285. doi:10.1016/0198-0149(87)90086-0

Menzel Barraqueta, J.-L., C. Sclosser, H. Planquette, A. Gouarin, M. Cheize, J. Boutorh, R. Shelley, L. Contreira Pereira, M. Gledhill, M.J. Hopwood, F. Lacan, P. Lherminier, G. Sarthou, and E. Achterberg. 2018. Aluminium in the North Atlantic Ocean and the Labrador Sea (GEOTRACES GA01 section): roles of continental inputs and biogenic particle removal. Biogeosciences 15: 5271–5286. doi:10.5194/bg-15-5271-2018

Nagata, T., Y. Yamada, and H. Fukuda. 2021. Transparent Exopolymer Particles in Deep Oceans: Synthesis and Future Challenges. Gels 7: 75. doi:10.3390/gels7030075

Nowicki, M., DeVries, T., & Siegel, D. A. (2022). Quantifying the Carbon Export and Sequestration Pathways of the Ocean’s Biological Carbon Pump. Global Biogeochemical Cycles, 36(3), e2021GB007083. 10.1029/2021GB007083

Omand, M. M., R. Govindarajan, J. He, and A. Mahadevan. 2020. Sinking flux of particulate organic matter in the oceans: Sensitivity to particle characteristics. Sci Rep 10: 5582. doi:10.1038/s41598-020-60424-5

Owens, S. A., S. Pike, and K. O. Buesseler. 2015. Thorium-234 as a tracer of particle dynamics and upper ocean export in the Atlantic Ocean. Deep Sea Research Part II: Topical Studies in Oceanography 116: 42–59. doi:10.1016/j.dsr2.2014.11.010

Passow, U. 2000. Formation of Transparent Exopolymer Particles, TEP, from dissolved precursor material. Marine ecology-progress series 192: 1–11.

Passow, U. 2002. Transparent exopolymer particles (TEP) in aquatic environments. Progress in Oceanography 55: 287–333. doi:10.1016/S0079-6611(02)00138-6

Passow, U. 2004. Switching perspectives: Do mineral fluxes determine particulate organic carbon fluxes or vice versa? Geochemistry, Geophysics, Geosystems 5. doi:10.1029/2003GC000670

Passow, U., and A. L. Alldredge. 1995. A dye-binding assay for the spectrophotometric measurement of transparent exopolymer particles (TEP). Limnology and Oceanography 40: 1326–1335. doi:10.4319/lo.1995.40.7.1326

Passow, U., A. L. Alldredge, and B. E. Logan. 1994. The role of particulate carbohydrate exudates in the flocculation of diatom blooms. Deep Sea Research Part I: Oceanographic Research Papers 41: 335–357. doi:10.1016/0967-0637(94)90007-8

Passow, U., R. F. Shipe, A. Murray, D. K. Pak, M. A. Brzezinski, and A. L. Alldredge. 2001. The origin of transparent exopolymer particles (TEP) and their role in the sedimentation of particulate matter. Continental Shelf Research 21: 327–346. doi:10.1016/S0278-4343(00)00101-1

Passow, U., and P. Wassmann. 1994. On the trophic fate of Phaeocystis pouchetii (Hariot): IV. The formation of marine snow by P. pouchetii. Marine Ecology Progress Series 104: 153–161.

Reigstad, M., and P. Wassmann. 2007. Does Phaeocystis spp. Contribute Significantly to Vertical Export of Organic Carbon? Biogeochemistry 83: 217–234.

Riebesell, U., L.T. Bach, R. G. J. Bellerby, J.F.R. Bermudez, T. Boxhammer, J. Czerry, A. Larzen, A. Ludwig, and K. G. Schulz. 2017. Competitive fitness of a predominant pelagic calcifier impaired by ocean acidification. Nature Geoscience 10: 19–23. doi:10.1038/ngeo2854

Riebesell, U., M. Reigstad, P. Wassmann, T. Noji, and U. Passow. 1995. On the trophic fate of *Phaeocystis pouchetii* (hariot): VI. Significance of *Phaeocystis*-derived mucus for vertical flux. Netherlands Journal of Sea Research 33: 193–203. doi:10.1016/0077-7579(95)90006-3

Riley, J. S., R. Sanders, C. Marsay, F. a. C. Le Moigne, E. P. Achterberg, and A. J. Poulton. 2012. The relative contribution of fast and slow sinking particles to ocean carbon export. Global Biogeochemical Cycles 26. doi:10.1029/2011GB004085

Romanelli, E., S. L. C. Giering, M. Estapa, D. A. Siegel, and U. Passow. 2024. Can intense storms affect sinking particle dynamics after the North Atlantic spring bloom? Limnology and Oceanography 69: 2963–2974. doi:10.1002/lno.12723

Romanelli, E., R. Stevens-Green, C. Cisternas-Novoa, J. LaRoche, D. A. Siegel, C. A. Carlson, and U. Passow. 2026. Particle lability drives degradation dynamics and bacterial community assembly during a Phaeocystis bloom decline. 2026.04.19.716305. doi:10.64898/2026.04.19.716305

Romanelli, E., J. Sweet, S. L. C. Giering, D. A. Siegel, and U. Passow. 2023. The importance of transparent exopolymer particles over ballast in determining both sinking and suspension of small particles during late summer in the Northeast Pacific Ocean. Elementa: Science of the Anthropocene 11: 00122. doi:10.1525/elementa.2022.00122

Rontani, J.-F., N. Zabeti, and S. G. Wakeham. 2011. Degradation of particulate organic matter in the equatorial Pacific Ocean: Biotic or abiotic? Limnology and Oceanography 56: 333–349. doi:10.4319/lo.2011.56.1.0333

Rousseau, V., M.-J. Chrétiennot-Dinet, A. Jacobsen, P. Verity, and S. Whipple. 2007. The Life Cycle of Phaeocystis: State of Knowledge and Presumptive Role in Ecology. Biogeochemistry 83: 29–47.

Sanders, R., S. A. Henson, M. Koski, C. L. De La Rocha, S. C. Painter, A. J. Poulton, J. Riley, B. Salihoglu, A. Visser, A. Yool, R. Bellerby, and A. P. Martin. 2014. The Biological Carbon Pump in the North Atlantic. Progress in Oceanography 129: 200–218. doi:10.1016/j.pocean.2014.05.005

Schoemann, V., S. Becquevort, J. Stefels, V. Rousseau, and C. Lancelot. 2005. Phaeocystis blooms in the global ocean and their controlling mechanisms: a review. Journal of Sea Research 53: 43–66. doi:10.1016/j.seares.2004.01.008

Siegel, D. A., A.B. Burd, M.L. Estapa, E. Fields, L. Johnson, U. Passow, E. Romanelli, M. A. Brzezinski, K. O. Busseler, S. J. Clevenger, I. Cetinic, L. Drago, C. A. Durkin, R. Kiko, S.J. Kramer, A. E. Maas, M. M. Omand, and D.K. Steinberg. 2025. Assessing Marine Snow Dynamics During the Demise of the North Atlantic Spring Bloom Using In Situ Particle Imagery. Global Biogeochemical Cycles 39: e2025GB008676. doi:10.1029/2025GB008676

Siegel, D. A., T. DeVries, I. Cetinić, and K. M. Bisson. 2023. Quantifying the Ocean’s Biological Pump and Its Carbon Cycle Impacts on Global Scales. Annual Review of Marine Science 15: 329–356. doi:10.1146/annurev-marine-040722-115226

Sieracki, C. K., M. E. Sieracki, and C. S. Yentsch. 1998. An imaging-in-flow system for automated analysis of marine microplankton. Marine Ecology Progress Series 168: 285–296. doi:10.3354/meps168285

Smetacek, V. 1985. Role of sinking in diatom life-history cycles: ecological, evolutionary and geological significance. Marine biology 84: 239–251.

Smith, W. O., D. J. McGillicuddy, E. B. Olson, V. Kosnyrev, E. E. Peacock, and H. M. Sosik. 2017. Mesoscale variability in intact and ghost colonies of *Phaeocystis antarctica* in the Ross Sea: Distribution and abundance. Journal of Marine Systems 166: 97–107. doi:10.1016/j.jmarsys.2016.05.007

Smith, W. O., and S. Trimborn. 2024. *Phaeocystis*: A Global Enigma. Annu. Rev. Mar. Sci. 16: annurev-marine-022223-025031. doi:10.1146/annurev-marine-022223-025031

Strickland, J. D. H., and T. R. Parsons. 1968. A Practical Handbook of Seawater Analyses.

Verity, P. G., C. P. Brussaard, J. C. Nejstgaard, M. A. van Leeuwe, C. Lancelot, and L. K. Medlin. 2007. Current understanding of Phaeocystis ecology and biogeochemistry, and perspectives for future research. Biogeochemistry 83: 311–330. doi:10.1007/s10533-007-9090-6

Verity, P. G., T. A. Villareal, and T. J. Smayda. 1988. Ecological investigations of blooms of colonial *Phaeocystis pouchetti*. II. The role of life-cycle phenomena in bloom termination. J Plankton Res 10: 749–766. doi:10.1093/plankt/10.4.749

Volk, T., and M. I. Hoffert. 1985. Ocean Carbon Pumps: Analysis of Relative Strengths and Efficiencies in Ocean-Driven Atmospheric CO2 Changes, p. 99–110. In The Carbon Cycle and Atmospheric CO2: Natural Variations Archean to Present. American Geophysical Union (AGU).

Wakeham, S. G., and C. Lee. 1993. Production, Transport, and Alteration of Particulate Organic Matter in the Marine Water Column, p. 145–169. In M.H. Engel and S.A. Macko [eds.], Organic Geochemistry: Principles and Applications. Springer US.

Der Wal, P. V., R. S. Kempers, and M. J. W. Veldhuis. 1995. Production and downward flux of organic matter and calcite in a North Sea bloom of the coccolithophore Emiliania huxleyi. Marine Ecology Progress Series 126: 247–265. doi:10.3354/meps126247

Wassmann, P., T. Ratkova, and M. Reigstad. 2005. The contribution of single and colonial cells of *Phaeocystis pouchetii* to spring and summer blooms in the north-eastern North Atlantic. Harmful Algae 4: 823–840. doi:10.1016/j.hal.2004.12.009

Williams, J. R., and S. L. C. Giering. 2022. In Situ Particle Measurements Deemphasize the Role of Size in Governing the Sinking Velocity of Marine Particles. Geophysical Research Letters 49: e2022GL099563. doi:10.1029/2022GL099563

Wolf, C., M. Iversen, C. Klaas, and K. Metfies. 2016. Limited sinking of Phaeocystis during a 12 days sediment trap study. Mol Ecol 25: 3428–3435. doi:10.1111/mec.13697

Wurl, O., and M. Holmes. 2008. The gelatinous nature of the sea-surface microlayer. Marine Chemistry 110: 89–97. doi:10.1016/j.marchem.2008.02.009

Yamada, Y., A. Ebihara, H. Fukuda, S. Otosaka, S. Mitarai, and T. Nagata. 2024. Functions of extracellular polymeric substances in partitioning suspended and sinking particles in the upper oceans of two open ocean systems. Limnology and Oceanography 69: 1101–1114. doi:10.1002/lno.12554

Yokogawa Fluid Imaging Technologies, Inc. 2023. VisualSpreadsheet 6: Calculations and particle property definitions.

