## Supplementary material for "Differences between suspended and sinking particles regulate carbon flux in the upper mesopelagic during a *Phaeocystis* Bloom": CisternasNovoa et al_supplemental material

**Supporting Information**
for

### Differences between suspended and sinking particles regulate carbon flux in the upper mesopelagic during a *Phaeocystis* Bloom

Cisternas-Novoa[a*], C., Romanelli, E[b]., and Uta Passow[a]

[a] Ocean Science Centre, Memorial University of Newfoundland, St. John's, NL, Canada

[b] Institute of Environmental Engineering, Department of Civil, Environmental and Geomatic Engineering, ETH Zurich, Switzerland

**Table S1. Calculation of Particle Concentration**

Calculations of concentrations for suspended and sinking particles ( Riley et al. 2012; Giering et al. 2016):

$$C_{sus} = C_{top} \quad (1)$$

$$C_{slow} = (C_{base} - C_{top})\ V_{\text{base}} / V_{\text{MSC}} \quad (2)$$

$$C_{fast} = (C_{tray} - C_{base})\ V_{\text{tray}} / (A_{\text{tray}}\ h_{\text{MSC}}) \quad (3)$$

With $V_{\text{base}}$ the volume of the base section (7.6 L), $V_{\text{MSC}}$ the volume of the MSC (89.9 L including base), $V_{\text{tray}}$ the volume of the tray (~1 L, measured for each deployment), $A_{\text{tray}}$ the area of the tray (0.026 $m^2$) and $h_{\text{MSC}}$ the height of the MSC (1.58 m).

**Table S2. POC normalized biogeochemical variables, also used for the Wilcoxon rank-sum test.** Uncertainties propagated assuming independent errors. TEP/POC (µg XG eq. (µg-C)$^{-1}$), CSP/POC (µg XG eq. (µg-C)$^{-1}$), C:N (molar), BSi/POC (µmol / µmol), and PIC/POC (µg/ µg)

| Bloom-stage | z | Particle type | TEP/POC | CSP/POC | C:N | BSi/POC | PICPOC |
|---|---|---|---|---|---|---|---|
| Late-bloom | D1 | sinking | 0.41 ± 0.23 | 0.69 ± 0.57 | 7.32 ± 1.39 | 0.01 | 0.20 |
| Late-bloom | D1 | suspended | 1.20 ± 0.07 | 0.39 ± 0.14 | 5.72 ± 0.42 | 0.01 | 0.07 |
| Late-bloom | D2 | sinking | 1.15 ± 0.03 | 0.88 ± 0.65 | 9.67 ± 1.48 | 0.01 | 0.03 |
| Late-bloom | D2 | suspended | 1.59 ± 0.31 | 0.42 ± 0.21 | 5.19 ± 1.17 | 0.01 | 0.02 |
| Late-bloom | D3 | sinking | 6.39 ± 8.68 | 2.77 ± 3.78 | 12.28 | 0.04 | 0.15 |
| Late-bloom | D3 | suspended | 3.58 ± 0.31 | 0.54 ± 0.51 | 5.20 ± 2.32 | 0.02 | 0.19 |
| Bloom-decline | D1 | sinking | 0.35 | 0.16 | 9.09 ± 2.80 | 0.01 | 0.00 |
| Bloom-decline | D1 | suspended | 1.30 | 0.46 | 5.77 ± 0.24 | 0.00 | 0.12 |
| Bloom-decline | D2 | sinking | 0.06 | 0.22 | 9.61 ± 1.12 | 0.02 | 0.00 |
| Bloom-decline | D2 | suspended | 6.12 | 0.55 | 5.13 ± 0.04 | 0.01 | 0.80 |
| Bloom-decline | D3 | sinking | 0.11 | 0.07 | 4.54 ± 4.76 | 0.01 | 0.00 |
| Bloom-decline | D3 | suspended | 2.37 | 0.49 | 4.91 ± 0.45 | 0.01 | 0.26 |
| Non-bloom | D1 | sinking | 0.14 | 0.24 | 9.81 ± 4.29 | 0.09 ± 0.00 | 0.40 ± 0.01 |
| Non-bloom | D1 | suspended | 0.90 | 0.15 | 4.78 ± 0.24 | 0.09 ± 0.01 | 0.11 ± 0.01 |
| Non-bloom | D2 | sinking | 0.09 | 0.08 | 7.31 ± 0.01 | 0.06 ± 0.01 | 0.27 ± 0.04 |
| Non-bloom | D2 | suspended | 1.01 | 0.05 | 5.03 ± 0.70 | 0.15 ± 0.01 | 0.72 ± 0.39 |
| Non-bloom | D3 | sinking | 0.09 | 0.14 | 9.89 ± 5.74 | 0.19± 0.01 | 0.44 ± 0.13 |
| Non-bloom | D3 | suspended | 2.11 | 0.19 | 4.80 ± 0.15 | 0.15 ± 0.03 | 0.07 ± 0.09 |

**Table S3. Variables used in the Factor Analysis of Mixed Data (FAMD).** Summary of average biogeochemical and morphological characteristics of suspended and sinking particles. Concentrations of exopolymeric particles and biominerals were normalized to particulate organic carbon (POC). TEP/POC (μg XG eq. $(\mu g\text{-}C)^{-1}$), CSP/POC (μg XG eq. $(\mu g\text{-}C)^{-1}$), C:N (molar), BSi (μmol / μmol), and PIC/POC (μg/μg). Morphological characteristics: area-based diameter (ABD, mm), slope of the PSD (particle size distribution), P*int* (a porosity proxy derived from intensity normalized by size), irregularity, and transparency were obtained using FlowCam.

| Categories | | | Biogeochemical composition | | | | | Morphological characteristics | | | | |
|---|---|---|---|---|---|---|---|---|---|---|---|---|
| Bloom stage | depth | particle type | TEP/ POC | CSP/ POC | BSi/ POC | PIC/ POC | C: N | ABD | Slope of PSD | $P_{int}$ | Irregularity | Trans-parency |
| Late-bloom | D1 | sinking | 0.45 | 0.78 | 0.01 | 0.20 | 7.0 | 24.9 | -2.0 | 8.4 | 1.28 | 0.16 |
| Late-bloom | D1 | suspended | 1.01 | 0.35 | 0.02 | 0.09 | 5.8 | 16.4 | -2.9 | 5.2 | 1.02 | 0.08 |
| Late-bloom | D2 | sinking | 1.06 | 1.16 | 0.01 | 0.04 | 6.6 | 17.9 | -2.3 | 6.2 | 1.11 | 0.13 |
| Late-bloom | D2 | suspended | 1.19 | 0.42 | 0.01 | 0.03 | 5.5 | 17.1 | -2.6 | 5.3 | 1.06 | 0.10 |
| Late-bloom | D3 | sinking | 5.48 | 4.63 | 0.04 | 0.14 | 10.4 | 17.6 | -2.0 | 5.7 | 1.25 | 0.13 |
| Late-bloom | D3 | suspended | 3.38 | 0.25 | 0.02 | 0.20 | 5.1 | 19.6 | -1.8 | 6.0 | 1.11 | 0.10 |
| Bloom-decline | D1 | sinking | 0.42 | 0.19 | 0.02 | 0.00 | 6.5 | 15.5 | -2.2 | 4.4 | 1.03 | 0.08 |
| Bloom-decline | D1 | suspended | 0.88 | 0.31 | 0.00 | 0.16 | 5.8 | 16.6 | -1.8 | 5.3 | 1.13 | 0.12 |
| Bloom-decline | D2 | sinking | 0.07 | 0.27 | 0.02 | 0.13 | 6.7 | 14.9 | -2.0 | 4.8 | 1.13 | 0.12 |
| Bloom-decline | D2 | suspended | 3.61 | 0.33 | 0.02 | 1.13 | 5.1 | 18.6 | -1.5 | 6.4 | 1.25 | 0.13 |
| Bloom-decline | D3 | sinking | 0.13 | 0.08 | 0.01 | 0.00 | 1.7 | 16.8 | -1.4 | 7.4 | 2.06 | 0.29 |
| Bloom-decline | D3 | suspended | 2.22 | 0.46 | 0.01 | 0.27 | 4.9 | 15.6 | -2.6 | 4.6 | 1.07 | 0.09 |
| Non-bloom | D1 | sinking | 0.46 | 0.46 | 0.12 | 0.61 | 5.8 | 30.4 | -1.2 | 12.9 | 2.31 | 0.18 |
| Non-bloom | D1 | suspended | 1.03 | 0.18 | 0.09 | 0.11 | 4.8 | 16.0 | -2.1 | 4.8 | 1.07 | 0.09 |
| Non-bloom | D2 | sinking | 0.14 | 0.14 | 0.07 | 0.28 | 1.8 | 18.9 | -1.6 | 5.6 | 1.13 | 0.10 |
| Non-bloom | D2 | suspended | 1.07 | 0.05 | 0.14 | 1.36 | 5.0 | 20.0 | -1.9 | 6.2 | 1.13 | 0.12 |
| Non-bloom | D3 | sinking | 0.19 | 0.30 | 0.10 | 0.53 | 5.6 | 18.1 | -2.0 | 6.0 | 1.12 | 0.11 |
| Non-bloom | D3 | suspended | 1.72 | 0.15 | 0.17 | 0.07 | 4.8 | 18.5 | -2.7 | 5.1 | 1.02 | 0.08 |

**Table S4. Particle Characteristics at D1 used to define the bloom-stage.** Values are presented as means ± standard deviations, except for the POC attenuation coefficient, which is shown as mean ± standard error. The pheopigments-to-Chl-a ratio includes both pheophorbide-a and pheophytin-a (Pheos). Statistical differences among bloom states (late-bloom: LB; bloom-decline: BD; and non-bloom: NB) were assessed using one-way ANOVA (significance threshold: $p < 0.05$), followed by Tukey's post-hoc tests to identify significant pairwise differences, which are indicated in the table.

| Parameters | Unit | Late-bloom (st. 24, 4, & 6) | Bloom-decline (st. 8 & 28) | Non-bloom (st. 15, 13, & 7) | Significant pairwise differences |
|---|---|---|---|---|---|
| Sinking marine snow concentration at D1 | ppm | 1102.2 ± 446.5 | 112.8 ± 56.1 | 18.3 ± 5.3 | BD vs LB; NB vs LB |
| Slope of sinking marine snow distribution at D1 | — | -1.63 ± 0.16 | -1.91 ± 0.37 | -2.29 ± 0.08 | NB vs LB |
| Sinking POC at D1 | mg/L | 28.77 ± 2.01 | 0.53 ± 0.25 | 0.32 ± 0.11 | NB vs LB |
| C:N ratio of sinking particles at D1 | molar | 7.33 ± 1.37 | 9.10 ± 2.83 | 9.81 ± 4.27 | None |
| Chl-a concentration at D1 | mg/L | 1.20 ± 0.33 | 0.19 ± 0.00 | 0.14 ± 0.04 | BD vs LB; NB vs LB |
| Pheos/Chl-a ratio at D1 | mg/mg | 0.05 ± 0.05 | 0.23 ± 0.05 | 0.76 ± 0.40 | NB vs LB |
| Instantaneous POC flux attenuation coefficient, *b* | — | 1.52 ± 0.29 | 0.67 ± 0.19 | 0.53 ± 0.02 | NB vs LB |

**Table S5. Results of nonparametric tests comparing particle morphological characteristics among all bloom stages** (late-bloom: LB, bloom-decline: BD, non-bloom: NB) for each particle type (sinking, suspended). For each parameter, area-based diameter (ABD), a proxy for particle porosity ($P_{Int}$), Irregularity, and transparency. The Kruskal–Wallis test reports the overall difference among bloom stages (p-value; effect size $\varepsilon^2 = H/(N-1)$). Pairwise differences were evaluated using Dunn's post-hoc test with Benjamini–Hochberg–adjusted p-values and effect sizes ($r = |Z|/\sqrt{N}$). Effect sizes are interpreted as small ($r < 0.1$), moderate (0.1–0.3), and large (> 0.3).

| Parameter | Particle Type | Test | Comparison | p-value | Effect size |
|---|---|---|---|---|---|
| ABD | sinking | Kruskal-Wallis | - | <0.001 | 0.01 |
| ABD | sinking | Dunn | LB- BD | <0.001 | 0.11 |
| ABD | sinking | Dunn | BD-NB | <0.001 | 0.01 |
| ABD | sinking | Dunn | DB - NB | <0.001 | 0.10 |
| ABD | suspended | Kruskal-Wallis | - | <0.001 | 0.00 |
| ABD | suspended | Dunn | LB -BD | <0.001 | 0.04 |
| ABD | suspended | Dunn | LB -NB | <0.001 | 0.04 |
| ABD | suspended | Dunn | DB- NB | <0.001 | 0.06 |
| $P_{int}$ | sinking | Kruskal-Wallis | - | <0.001 | 0.02 |
| $P_{int}$ | sinking | Dunn | LB - BD | <0.001 | 0.15 |
| $P_{int}$ | sinking | Dunn | LB -NB | <0.001 | 0.06 |
| $P_{int}$ | sinking | Dunn | BD - NB | <0.001 | 0.10 |
| $P_{int}$ | suspended | Kruskal-Wallis | - | <0.001 | 0.00 |
| $P_{int}$ | suspended | Dunn | LB - BD | <0.001 | 0.04 |
| $P_{int}$ | suspended | Dunn | LB - NB | <0.001 | 0.02 |
| $P_{int}$ | suspended | Dunn | BD - NB | <0.001 | 0.02 |
| Irregularity | sinking | Kruskal-Wallis | - | <0.001 | 0.01 |
| Irregularity | sinking | Dunn | LB - BD | <0.001 | 0.08 |
| Irregularity | sinking | Dunn | LB - NB | <0.001 | 0.06 |
| Irregularity | sinking | Dunn | BD- NB | <0.001 | 0.04 |
| Irregularity | suspended | Kruskal-Wallis | - | <0.001 | 0.00 |
| Irregularity | suspended | Dunn | LB - BD | <0.001 | 0.06 |
| Irregularity | suspended | Dunn | LB- NB | <0.01 | 0.01 |
| Irregularity | suspended | Dunn | BD - NB | <0.001 | 0.04 |
| Transparency | sinking | Kruskal-Wallis | - | <0.001 | 0.01 |
| Transparency | sinking | Dunn | LB - BD | <0.001 | 0.08 |
| Transparency | sinking | Dunn | LB - NB | <0.001 | 0.05 |

| | | | | | |
|---|---|---|---|---|---|
| Transparency | sinking | Dunn | BD - NB | <0.001 | 0.04 |
| Transparency | suspended | Kruskal-Wallis | - | <0.001 | 0.00 |
| Transparency | suspended | Dunn | LB- BD | <0.001 | 0.05 |
| Transparency | suspended | Dunn | LB - NB | 0.55 | 0.00 |
| Transparency | suspended | Dunn | BD - NB | <0.001 | 0.04 |

**Table S6. Wilcoxon rank-sum test results** comparing biogeochemical variables between suspended and sinking particles across all bloom states and depths. Differences were statistically significant for C:N and TEP/POC ($p \leq 0.01$, in bold).

| Variable | n | p_value |
|---|---|---|
| **TEP/POC** | **18** | **0.010** |
| CSP/POC | 18 | 0.863 |
| **C:N** | **18** | **0.004** |
| BSi/POC | 18 | 0.711 |
| PIC/POC | 18 | 0.535 |

**Table S7. Particle morphological characteristics.** Presented for each bloom stage, depth, and particle type. Sample size (n); mean ± SD of area-based diameter (ABD, mm); slope and $r^2$ of the particle size distribution (PSD); and mean ± SD of porosity proxy ($P_{int}$), irregularity, and transparency.

| Bloom stage | depth | particle type | n | ABD | PSD slope | slope r2 | $P_{int}$ | Irregu- larity | Trans- parency |
|---|---|---|---|---|---|---|---|---|---|
| Late-bloom | D1 | sinking | 75096 | 24.9± 19.0 | -1.96 | 0.89 | 8.4 ± 7.2 | 1.3 ± 0.7 | 0.16 ± 0.14 |
| Late-bloom | D1 | suspended | 10084 0 | 16.4 ± 5.7 | -2.90 | 0.97 | 5.2 ± 2.6 | 1.0 ± 0.1 | 0.08 ± 0.06 |
| Late-bloom | D2 | sinking | 89796 | 17.9 ± 9.9 | -2.34 | 0.96 | 6.2 ± 5.3 | 1.1± 0.4 | 0.13 ± 0.15 |
| Late-bloom | D2 | suspended | 54233 | 17.1 ± 8.5 | -2.62 | 0.97 | 5.3 ± 3.3 | 1.1 ± 0.5 | 0.10 ± 0.12 |
| Late-bloom | D3 | sinking | 5900 | 17.6 ± 14.3 | -1.96 | 0.98 | 5.7 ± 5.3 | 1.3 ± 0.8 | 0.13 ± 0.14 |
| Late-bloom | D3 | suspended | 8531 | 19.6 ± 12.0 | -1.79 | 0.95 | 6.0 ± 6.1 | 1.1 ± 0.5 | 0.10 ± 0.10 |
| Bloom-decline | D1 | sinking | 6913 | 15.5± 7.8 | -2.20 | 0.96 | 4.4 ± 3.1 | 1.0 ± 0.3 | 0.08 ± 0.05 |
| Bloom-decline | D1 | suspended | 2900 | 16.6 ± 9.7 | -1.81 | 0.98 | 5.3 ± 3.5 | 1.1 ± 0.4 | 0.12 ± 0.10 |
| Bloom-decline | D2 | sinking | 2240 | 14.9 ± 8.2 | -2.01 | 0.96 | 4.8 ± 3.8 | 1.1 ± 0.4 | 0.12 ± 0.14 |
| Bloom-decline | D2 | suspended | 4265 | 18.6 ± 16.2 | -1.50 | 0.99 | 6.4 ± 7.9 | 1.3 ± 0.8 | 0.13 ± 0.14 |
| Bloom-decline | D3 | sinking | 420 | 16.8 ± 15.2 | -1.36 | 0.97 | 7.4 ± 9.1 | 2.1 ± 1.6 | 0.29 ± 0.26 |
| Bloom-decline | D3 | suspended | 5204 | 15.6 ± 6.6 | -2.57 | 0.97 | 4.6 ± 2.5 | 1.1 ± 0.3 | 0.09 ± 0.07 |
| Non-bloom | D1 | sinking | 9575 | 30.4 ± 31.7 | -1.23 | 0.88 | 12.9 ± 17.7 | 2.3 ± 2.9 | 0.18 ± 0.16 |
| Non-bloom | D1 | suspended | 3643 | 16.0 ± 8.8 | -2.11 | 0.95 | 4.8 ± 4.5 | 1.1 ± 0.4 | 0.09 ± 0.09 |
| Non-bloom | D2 | sinking | 4506 | 18.9 ± 13.7 | -1.57 | 0.97 | 5.7 ± 6.5 | 1.1 ± 0.8 | 0.10 ± 0.10 |
| Non-bloom | D2 | suspended | 2035 | 19.9 ± 11.5 | -1.94 | 0.94 | 6.2 ± 4.7 | 1.1 ± 0.4 | 0.12 ± 0.10 |
| Non-bloom | D3 | sinking | 18184 | 18.1 ± 10.0 | -1.98 | 0.98 | 6.0 ± 5.6 | 1.1 ± 0.6 | 0.11 ± 0.13 |
| Non-bloom | D3 | suspended | 6406 | 18.5± 7.3 | -2.70 | 0.92 | 5.1 ± 2.5 | 1.0 ± 0.1 | 0.08 ± 0.06 |

**Table S8. Sinking marine snow size and concentration at D1 (shallow MSC).** Mean ± standard deviation of particle equivalent spherical diameter (ESD, mm), concentration (# $L^{-1}$), and total volume (ppm), quantified from images of MSC trays.

| Bloom-stage | ESD (mm) | Concentration (# $L^{-1}$) | Concentration (ppm) |
|---|---|---|---|
| Late-bloom | 0.28 ± 0.24 | 6416 ± 2181 | 763 ± 155 |
| Bloom-decline | 0.25 ± 0.16 | 4320 ± 4965 | 112 ± 20 |
| Non-bloom | 0.21 ± 0.18 | 1397 ± 168 | 68 ± 4 |

**Table S9. Summary of sinking marine snow from visual inspection of MSC trays.** Marine snow amount, size, and appearance using relative measures. NA: no analysis.

| Bloom-stage | station | depth | MS amount | MS size | MS characteristics |
|---|---|---|---|---|---|
| **Late-bloom** | 24 | D1 | moderate | medium | brown |
| **Late-bloom** | 4 | D1 | high | medium | green |
| **Late-bloom** | 6 | D1 | high | large | green, fluffy, and fragile |
| **Late-bloom** | 24 | D2 | moderate | small | round and dark |
| **Late-bloom** | 4 | D2 | moderate | NA | compact |
| **Late-bloom** | 6 | D2 | moderate | NA | NA |
| **Late-bloom** | 24 | D3 | moderate | small | compact, 2 small fecal pellets (FP) |
| **Late-bloom** | 4 | D3 | minimal | large | compact |
| **Late-bloom** | 6 | D3 | low | small | compact |
| **Bloom-decline** | 8 | D1 | moderate | all sizes | green |
| **Bloom-decline** | 28 | D1 | moderate | small and few Large | green, fluffy |
| **Bloom-decline** | 8 | D2 | moderate | medium and small | NA |
| **Bloom-decline** | 28 | D2 | moderate | small | fluffy |
| **Bloom-decline** | 8 | D3 | low | small | compact |
| **Bloom-decline** | 28 | D3 | low | small | compact |
| **Non-bloom** | 15 | D1 | low | small | round, compact |
| **Non-bloom** | 13 | D1 | low | small and medium | fluffy (medium size) |
| **Non-bloom** | 7 | D1 | low | small and Large | fluffy |
| **Non-bloom** | 15 | D2 | low | few medium, mostly small | NA |
| **Non-bloom** | 13 | D2 | low | large and small | fluffy- white (Large), compact (smaller) |
| **Non-bloom** | 7 | D2 | low | large and small | fluffy- green (Large), compact (smaller) |
| **Non-bloom** | 15 | D3 | low | medium and small | NA |
| **Non-bloom** | 13 | D3 | low | large and small | fluffy (Large), compact and white (small), and 1 round, compact and dark (Large) |
| **Non-bloom** | 7 | D3 | low | large and small | 2FP (Large), compact and green (Large), white mixed compact and fluffy (small) |

**Table S10. Variable contributions to the first five dimensions of the Factor Analysis of Mixed Data (FAMD).** Contributions > 0.5 (green) and 0.15–0.5 (blue) are highlighted to indicate strong and moderate associations with each dimension. Discussion of dimensions 1 and 2 is provided in the main text. Dimension 3 showed the highest contributions from the bloom stage (0.58) and BSi/POC (0.47), with smaller contributions from transparency (0.19) and TEP/POC (0.17), capturing siliceous and optical variations associated with the bloom stage. Dimensions 4 and 5 were primarily influenced by depth, with contributions of 0.57 and 0.69, respectively, and dimension 4 also included moderate contributions from TEP/POC (0.28) and PIC/POC (0.24), suggesting vertical structuring related to particle remineralization and carbonate content. The categorical variable "particle type" (suspended vs. sinking) contributed only modestly to the construction of the axes (maximum = 0.19 on Dimension 1), highlighting a key strength of FAMD: Even when a categorical variable does not strongly influence the dimensions, clear group separation can still emerge if it aligns with underlying quantitative trait patterns. The observed separation stems not from the label particle type itself, but from distinct trait profiles associated with suspended and sinking particles.

| Functional Clarification | Variable | Dim.1 28% | Dim.2 19% | Dim.3 14% | Dim.4 11% | Dim.5 9% |
|---|---|---|---|---|---|---|
| Environmental context | bloom-stage | 0.19 | 0.47 | 0.58 | 0.15 | 0.10 |
| | depth | 0.02 | 0.01 | 0.01 | 0.58 | 0.69 |
| Particle type | suspended/sinking | 0.19 | 0.17 | 0.06 | 0.03 | 0.02 |
| Exopolymeric particles | TEP/POC | 0.12 | 0.30 | 0.17 | 0.28 | 0.00 |
| | CSP/POC | 0.04 | 0.71 | 0.10 | 0.02 | 0.01 |
| Organic matter quality | C:N | 0.10 | 0.47 | 0.13 | 0.04 | 0.04 |
| Biominerals | BSi/POC | 0.13 | 0.20 | 0.47 | 0.00 | 0.12 |
| | PIC/POC | 0.13 | 0.10 | 0.21 | 0.24 | 0.22 |
| Morphological characteristics | area-based diameter (ABD) | 0.61 | 0.04 | 0.16 | 0.06 | 0.02 |
| | PSD slope | 0.60 | 0.02 | 0.02 | 0.13 | 0.02 |
| | irregularity | 0.78 | 0.07 | 0.03 | 0.002 | 0.12 |
| | transparency | 0.49 | 0.10 | 0.19 | 0.05 | 0.01 |
| | $P_{int}$ (porosity proxy) | 0.8 | 0.07 | 0.03 | 0.03 | 0.02 |

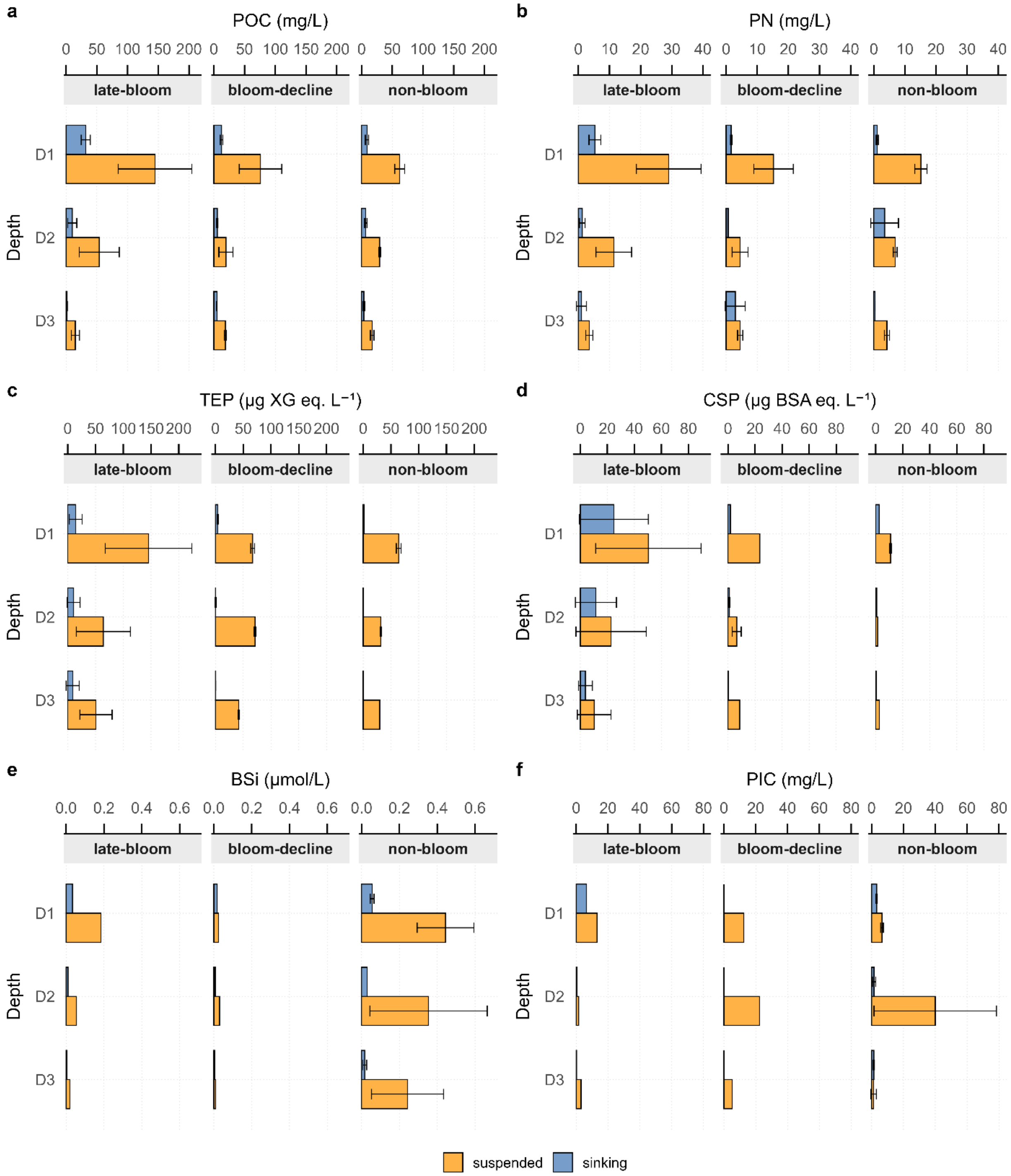


**Figure S1. Particle composition for all bloom-stages, depths, and particle types.** Concentrations of organic components (POC, PN, TEP, CSP) and biominerals (BSi, PIC) for all bloom-stages, depths (D1:70–150 m; D2: 125–250 m; D3: 300–500 m), and particle types (suspended: yellow; sinking: blue). Error bars indicate ± standard deviation.

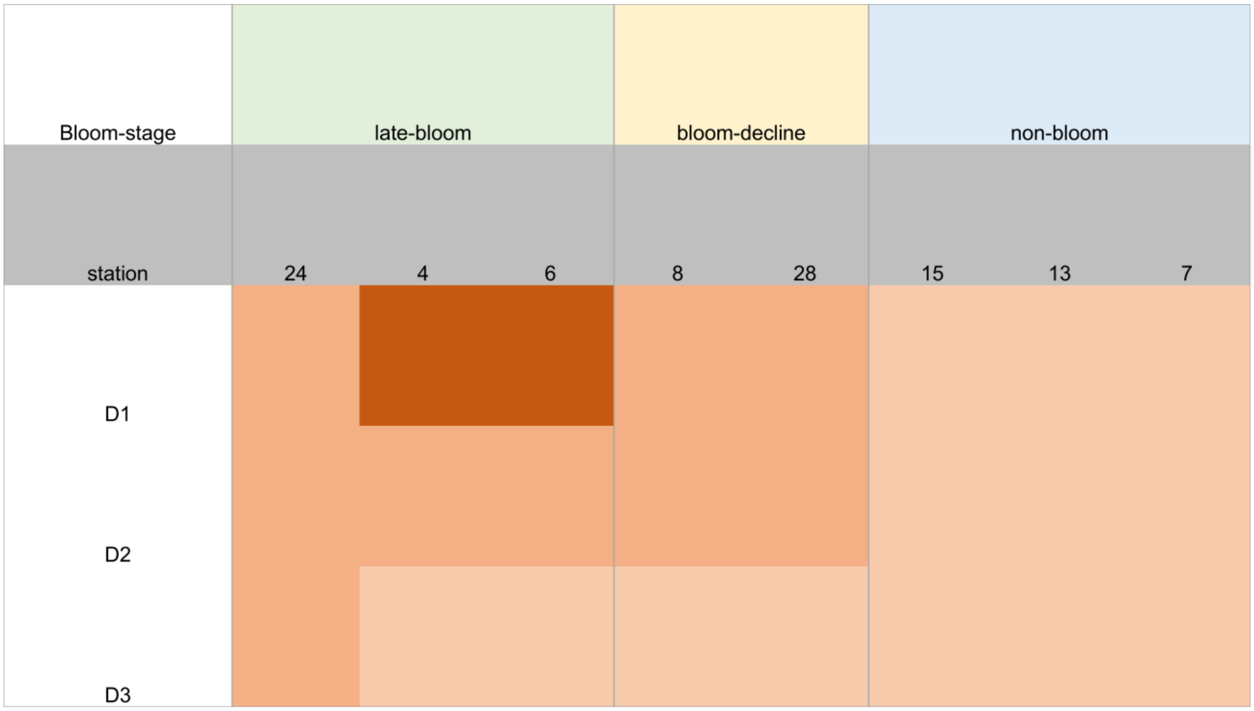


**Figure S2. Heat map showing the relative abundance of sinking marine snow based on visual inspection of MSC trays.** The stations of each bloom stage and the MS amount at each depth are depicted: high (dark orange), moderate (medium orange), low (light orange), and single aggregate (lightest orange).

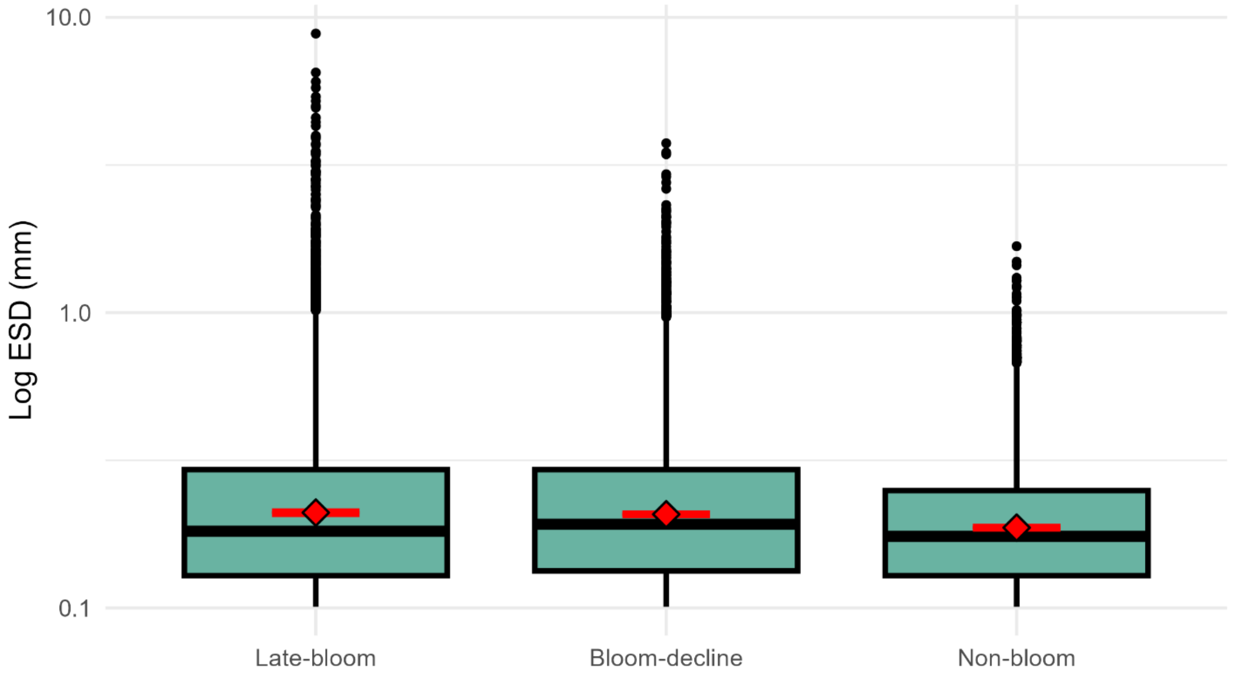


**Figure S3. Size of sinking marine snow (>0.1 mm), given as equivalent spherical diameter (ESD) at D1**. Boxes show the median and interquartile range, with outliers shown as points. Red points indicate the mean. A logarithmic scale is used to capture the full range of particle sizes, including extreme values.
